# Analysis of protein-coding genetic variation in 60,706 humans

**DOI:** 10.1101/030338

**Authors:** Exome Aggregation Consortium, Monkol Lek, Konrad J Karczewski, Eric V Minikel, Kaitlin E Samocha, Eric Banks, Timothy Fennell, Anne H O’Donnell-Luria, James S Ware, Andrew J Hill, Beryl B Cummings, Taru Tukiainen, Daniel P Birnbaum, Jack A Kosmicki, Laramie E Duncan, Karol Estrada, Fengmei Zhao, James Zou, Emma Pierce-Hoffman, Joanne Berghout, David N Cooper, Nicole Deflaux, Mark DePristo, Ron Do, Jason Flannick, Menachem Fromer, Laura Gauthier, Jackie Goldstein, Namrata Gupta, Daniel Howrigan, Adam Kiezun, Mitja I Kurki, Ami Levy Moonshine, Pradeep Natarajan, Lorena Orozco, Gina M Peloso, Ryan Poplin, Manuel A Rivas, Valentin Ruano-Rubio, Samuel A Rose, Douglas M Ruderfer, Khalid Shakir, Peter D Stenson, Christine Stevens, Brett P Thomas, Grace Tiao, Maria T Tusie-Luna, Ben Weisburd, Hong-Hee Won, Dongmei Yu, David M Altshuler, Diego Ardissino, Michael Boehnke, John Danesh, Stacey Donnelly, Roberto Elosua, Jose C Florez, Stacey B Gabriel, Gad Getz, Stephen J Glatt, Christina M Hultman, Sekar Kathiresan, Markku Laakso, Steven McCarroll, Mark I McCarthy, Dermot McGovern, Ruth McPherson, Benjamin M Neale, Aarno Palotie, Shaun M Purcell, Danish Saleheen, Jeremiah M Scharf, Pamela Sklar, Patrick F Sullivan, Jaakko Tuomilehto, Ming T Tsuang, Hugh C Watkins, James G Wilson, Mark J Daly, Daniel G MacArthur

## Abstract

Large-scale reference data sets of human genetic variation are critical for the medical and functional interpretation of DNA sequence changes. Here we describe the aggregation and analysis of high-quality exome (protein-coding region) sequence data for 60,706 individuals of diverse ethnicities generated as part of the Exome Aggregation Consortium (ExAC). The resulting catalogue of human genetic diversity contains an average of one variant every eight bases of the exome, and provides direct evidence for the presence of widespread mutational recurrence. We show that this catalogue can be used to calculate objective metrics of pathogenicity for sequence variants, and to identify genes subject to strong selection against various classes of mutation; we identify 3,230 genes with near-complete depletion of truncating variants, 72% of which have no currently established human disease phenotype. Finally, we demonstrate that these data can be used for the efficient filtering of candidate disease-causing variants, and for the discovery of human “knockout” variants in protein-coding genes.

## Background

Over the last five years, the widespread availability of high-throughput DNA sequencing technologies has permitted the sequencing of the whole genomes or exomes (the protein-coding regions of genomes) of hundreds of thousands of humans. In theory, these data represent a powerful source of information about the global patterns of human genetic variation, but in practice, are difficult to access for practical, logistical, and ethical reasons; in addition, their utility is complicated by the heterogeneity in the experimental methodologies and variant calling pipelines used to generate them. Current publicly available datasets of human DNA sequence variation contain only a small fraction of all sequenced samples: the Exome Variant Server, created as part of the NHLBI Exome Sequencing Project (ESP)^1^, contains frequency information spanning 6,503 exomes; and the 1000 Genomes (1000G) Project, which includes individual-level genotype data from whole-genome and exome sequence data for 2,504 individuals^2^.

Databases of genetic variation are important for our understanding of human population history and biology^1–5^, but also provide critical resources for the clinical interpretation of variants observed in patients suffering from rare Mendelian diseases^6,7^. The filtering of candidate variants by frequency in unselected individuals is a key step in any pipeline for the discovery of causal variants in Mendelian disease patients, and the efficacy of such filtering depends on both the size and the ancestral diversity of the available reference data.

Here, we describe the joint variant calling and analysis of high-quality variant calls across 60,706 human exomes, assembled by the Exome Aggregation Consortium (ExAC; exac.broadinstitute.org). This call set exceeds previously available exome-wide variant databases by nearly an order of magnitude, providing substantially increased resolution for the analysis of very low-frequency genetic variants. We demonstrate the application of this data set to the analysis of patterns of genetic variation including the discovery of widespread mutational recurrence, the inference of gene-level constraint against truncating variation, the clinical interpretation of variation in Mendelian disease genes, and the discovery of human “knockout” variants in protein-coding genes.

## The ExAC Data set

Sequencing data processing, variant calling, quality control and filtering was performed on over 91,000 exomes (see Online Methods), and sample filtering was performed to produce a final data set spanning 60,706 individuals (Figure 1a). To identify the ancestry of each ExAC individual, we performed principal component analysis (PCA) to distinguish the major axes of geographic ancestry and to identify population clusters corresponding to individuals of European, African, South Asian, East Asian, and admixed American (hereafter Latino) ancestry (Figure 1b; Supplementary Information Table 3); we note that the apparent separation between East Asian and other samples reflects a deficiency of Middle Eastern and Central Asian samples in the data set. We further separated Europeans into individuals of Finnish and non-Finnish ancestry given the enrichment of this bottlenecked population; the term “European” hereafter refers to non-Finnish European individuals.

**Figure 1.**
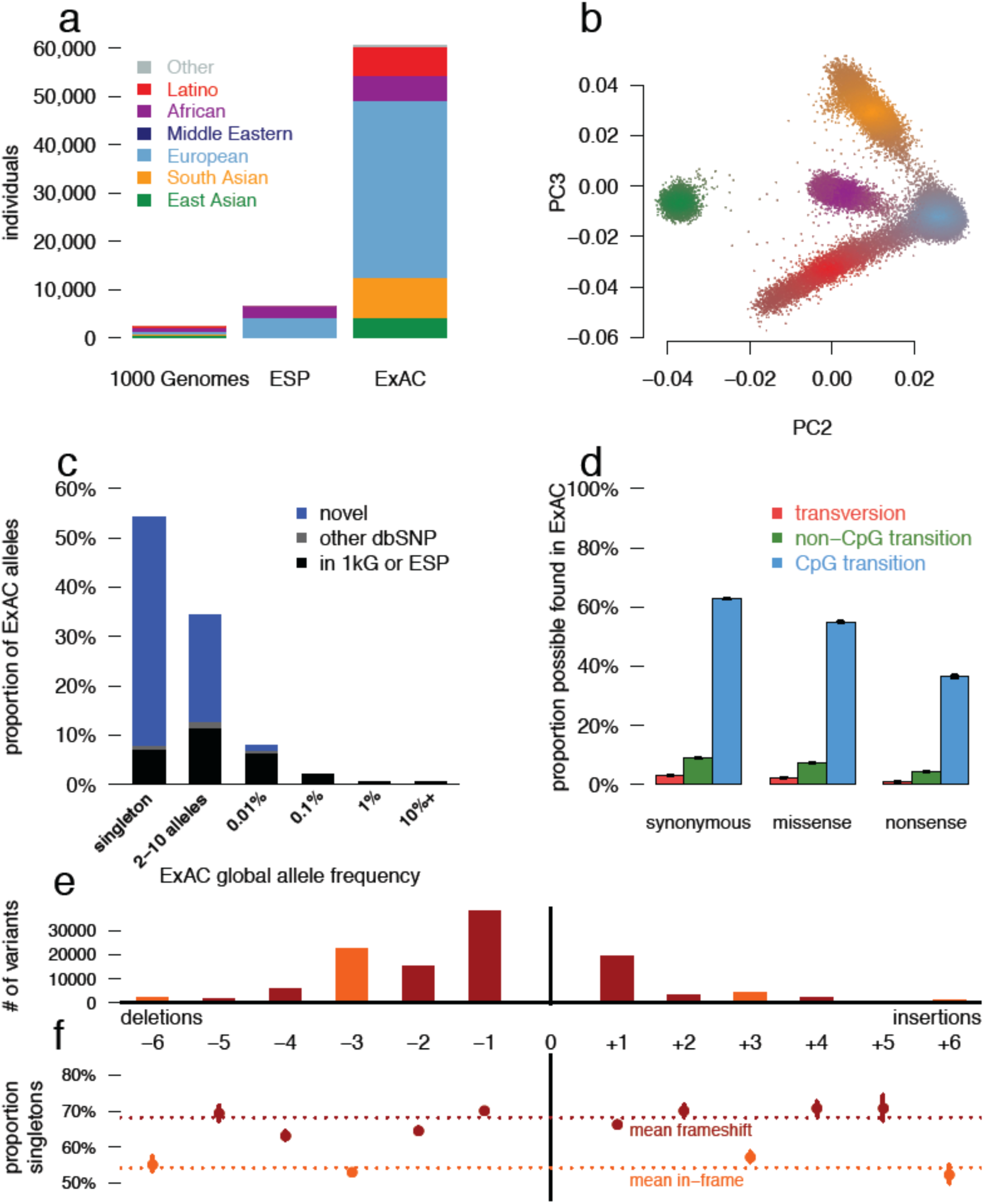
Patterns of genetic variation in 60,706 humans. a) The size and diversity of public reference exome datasets. ExAC exceeds previous datasets in size for all studied populations. b) Principal component analysis (PCA) dividing ExAC individuals into five continental populations. PC2 and PC3 are shown; additional PCs are in Extended Data Figure 2a. c) The allele frequency spectrum of ExAC highlights that the majority of genetic variants are rare and novel. d) The proportion of possible variation observed by mutational context and functional class. Over half of all possible CpG transitions are observed. Error bars represent standard error of the mean. e-f) The number (e) and frequency distribution (proportion singleton; f) of indels, by size. Compared to in-frame indels, frameshift variants are less common (have a higher proportion of singletons, a proxy for predicted deleteriousness on gene product). Error bars indicate 95% confidence intervals.

We identified 10,195,872 candidate sequence variants in ExAC. We further applied stringent depth and site/genotype quality filters to define a subset of 7,404,909 high quality (HQ) variants, including 317,381 indels (Supplementary Information Table 7), corresponding to one variant for every 8 bp within the exome intervals. The majority of these are very low-frequency variants absent from previous smaller call sets (Figure 1c): of the HQ variants, 99% have a frequency of <1%, 54% are singletons (variants seen only once in the data set), and 72% are absent from both 1000G and ESP.

The density of variation in ExAC is not uniform across the genome, and the observation of variants depends on factors such as mutational properties and selective pressures. In the ~45M well covered (80% of individuals with a minimum of 10X coverage) positions in ExAC, there are ~18M possible synonymous variants, of which we observe 1.4M (7.5%). However, we observe 63.1% of possible CpG transitions (C to T variants, where the adjacent base is G), while only observing 3% of possible transversions and 9.2% of other possible transitions (Supplementary Information Table 9). A similar pattern is observed for missense and nonsense variants, with lower proportions due to selective pressures (Figure 1D). Of 123,629 HQ insertion/deletions (indels) called in coding exons, 117,242 (95%) have length <6 bases, with shorter deletions being the most common (Figure 1E). Frameshifts are found in smaller numbers and are more likely to be singletons than in-frame indels (Figure 1F), reflecting the influence of purifying selection.

## Patterns of protein-coding variation revealed by large samples

The density of protein-coding sequence variation in ExAC reveals a number of properties of human genetic variation undetectable in smaller data sets. For instance, 7.9% of HQ sites in ExAC are multiallelic (multiple different sequence variants observed at the same site), close to the Poisson expectation of 8.3% given the observed density of variation, and far higher than observed in previous data sets - 0.48% in 1000 Genomes (exome intervals) and 0.43% in ESP.

The size of ExAC also makes it possible to directly observe mutational recurrence: instances in which the same mutation has occurred multiple times independently throughout the history of the sequenced populations. For instance, among synonymous variants, a class of variation expected to have undergone minimal selection, 43% of validated *de novo* events identified in external datasets of 1,756 parent-offspring trios^8,9^ are also observed independently in our dataset (Figure 2a), indicating a separate origin for the same variant within the demographic history of the two samples. This proportion is much higher for transition variants at CpG sites, well established to be the most highly mutable sites in the human genome^10^: 87% of previously reported *de novo* CpG transitions at synonymous sites are observed in ExAC, indicating that our sample sizes are beginning to approach saturation of this class of variation. This saturation is detectable by a change in the discovery rate at subsets of the ExAC data set, beginning at around 20,000 individuals (Figure 2b), indicating that ExAC is the first human exome-wide dataset large enough for this effect to be directly observed.

**Figure 2.**
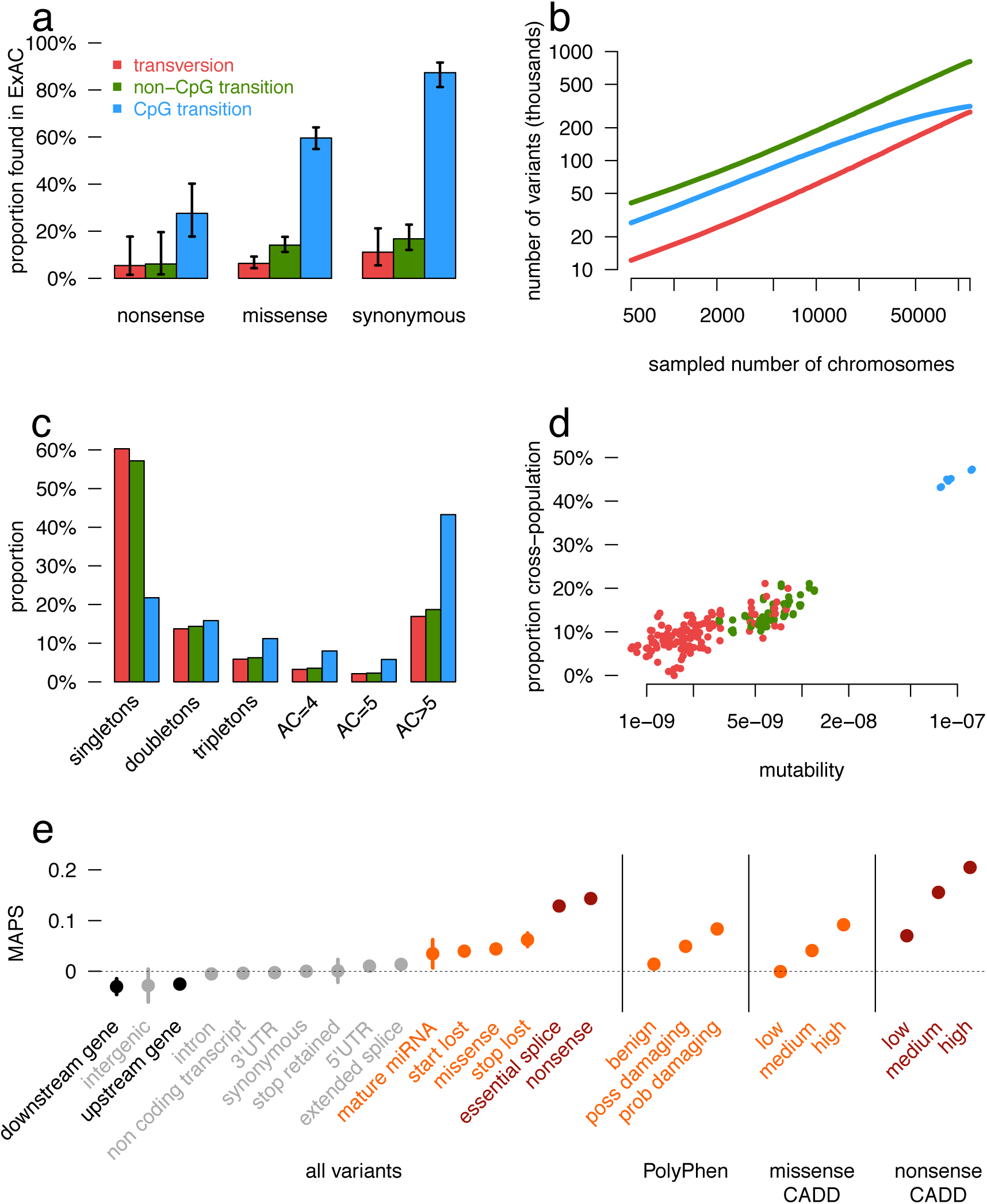
Mutational recurrence at large sample sizes. a) Proportion of validated *de novo* variants from two external datasets that are independently found in ExAC, separated by functional class and mutational context. Error bars represent standard error of the mean. Colors are consistent in a-d. b) Number of unique variants observed, by mutational context, as a function of number of individuals (down-sampled from ExAC). CpG transitions, the most likely mutational event, begin reaching saturation at ~20,000 individuals. c) The site frequency spectrum is shown for each mutational context. d) For doubletons (variants with an allele count of 2), mutation rate is positively correlated with the likelihood of being found in two individuals of different continental populations. e) The mutability-adjusted proportion of singletons (MAPS) is shown across functional classes. Error bars represent standard error of the mean of the proportion of singletons.

Mutational recurrence has a marked effect on the frequency spectrum in the ExAC data, resulting in a depletion of singletons at sites with high mutation rates (Figure 2c). We observe a correlation between singleton rates (the proportion of variants seen only once in ExAC) and site mutability inferred from sequence context^11^ (r = −0.98; p < 10^−50^; Extended Data Figure 4d): sites with low predicted mutability have a singleton rate of 60%, compared to 20% for sites with the highest predicted rate (CpG transitions; Figure 2C). Conversely, for synonymous variants, CpG variants are approximately twice as likely to rise to intermediate frequencies: 16% of CpG variants are found in at least 20 copies in ExAC, compared to 8% of transversions and non-CpG transitions, suggesting that synonymous CpG transitions have on average two independent mutational origins in the ExAC sample. Recurrence at highly mutable sites can further be observed by examining the population sharing of doubleton synonymous variants (variants occurring in only two individuals in ExAC). Low-mutability mutations (especially transversions), are more likely to be observed in a single population (representing a single mutational origin), while CpG transitions are more likely to be found in two separate populations (independent mutational events); as such, site mutability and probability of observation in two populations is significantly correlated (r = 0.884; Figure 2d).

We also explored the prevalence and functional impact of multinucleotide polymorphisms (MNPs), in cases where multiple substitutions were observed within the same codon in at least one individual. We found 5,945 MNPs (mean: 23 per sample) in ExAC (Extended Data Figure 3a) where analysis of the underlying SNPs without correct haplotype phasing would result in altered interpretation. These include 647 instances where the effect of a protein-truncating variant (PTV) variant is eliminated by an adjacent SNP (rescued PTV) and 131 instances where underlying synonymous or missense variants result in PTV MNPs (gained PTV). Additionally our analysis revealed 8 MNPs in disease-associated genes, resulting in either a rescued or gained PTV, and 10 MNPs that have previously been reported as disease causing mutations (Supplementary Information Table 10 and 11). We note that these variants would be missed by virtually all currently available variant calling and annotation pipelines.

## Inferring variant deleteriousness and gene constraint

Deleterious variants are expected to have lower allele frequencies than neutral ones, due to negative selection. This theoretical property has been demonstrated previously in human population sequencing data^12,13^ and here (Figure 1d, Figure 1e). This allows inference of the degree of selection against specific functional classes of variation: however, mutational recurrence as described above indicates that allele frequencies observed in ExAC-scale samples are also skewed by mutation rate, with more mutable sites less likely to be singletons (Figure 2c and Extended Data Figure 4d). Mutation rate is in turn non-uniformly distributed across functional classes - for instance, stop lost mutations can never occur at CpG dinucleotides (Extended Data Figure 4e). We corrected for mutation rates (Supplementary Information Section 3.2) by creating a mutability-adjusted proportion singleton (MAPS) metric. This metric reflects (as expected) strong selection against predicted PTVs, as well as missense variants predicted by conservation-based methods to be deleterious (Figure 2e).

The deep ascertainment of rare variation in ExAC also allows us to infer the extent of selection against variant categories on a per-gene basis by examining the proportion of variation that is missing compared to expectations under random mutation. Conceptually similar approaches have been applied to smaller exome datasets^11,14^ but have been underpowered, particularly when analyzing the depletion of PTVs. We compared the observed number of rare (MAF <0.1%) variants per gene to an expected number derived from a selection neutral, sequence-context based mutational model^11^. The model performs well in predicting the number of synonymous variants, which should be under minimal selection, per gene (r = 0.98; Extended Data Figure 5b).

We quantified deviation from expectation with a Z score^11^, which for synonymous variants is centered at zero, but is significantly shifted towards higher values (greater constraint) for both missense and PTV (Wilcoxon p < 10^−50^ for both; Figure 3a). The genes on the X chromosome are significantly more constrained than those on the autosomes for missense (p < 10^−7^) and loss-of-function (p < 10^−50^). The high correlation between the observed and expected number of synonymous variants on the X chromosome (r = 0.97 vs 0.98 for autosomes) indicates that this difference in constraint is not due to a calibration issue. To reduce confounding by coding sequence length for PTVs, we developed an expectation-maximization algorithm (Supplementary Information Section 4.4) using the observed and expected PTV counts within each gene to separate genes into three categories: null (observed ≈ expected), recessive (observed ≤50% of expected), and haploinsufficient (observed <10% of expected). This metric – the probability of being loss-of-function (LoF) intolerant (pLI) – separates genes of sufficient length into LoF intolerant (pLI ≥0.9, n=3,230) or LoF tolerant (pLI ≤0.1, n=10,374) categories. pLI is less correlated with coding sequence length (r = 0.17 as compared to 0.57 for the PTV Z score), outperforms the PTV Z score as an intolerance metric (Supplementary Information Table 15), and reveals the expected contrast between gene lists (Figure 3b). pLI is positively correlated with a gene product’s number of physical interaction partners (p < 10^−41^). The most constrained pathways (highest median pLI for the genes in the pathway) are core biological processes (spliceosome, ribosome, and proteasome components; KS test p < 10^−6^ for all) while olfactory receptors are among the least constrained pathways (KS test p < 10^−16^), demonstrated in Figure 3b and consistent with previous work^5,15–18^.

**Figure 3.**
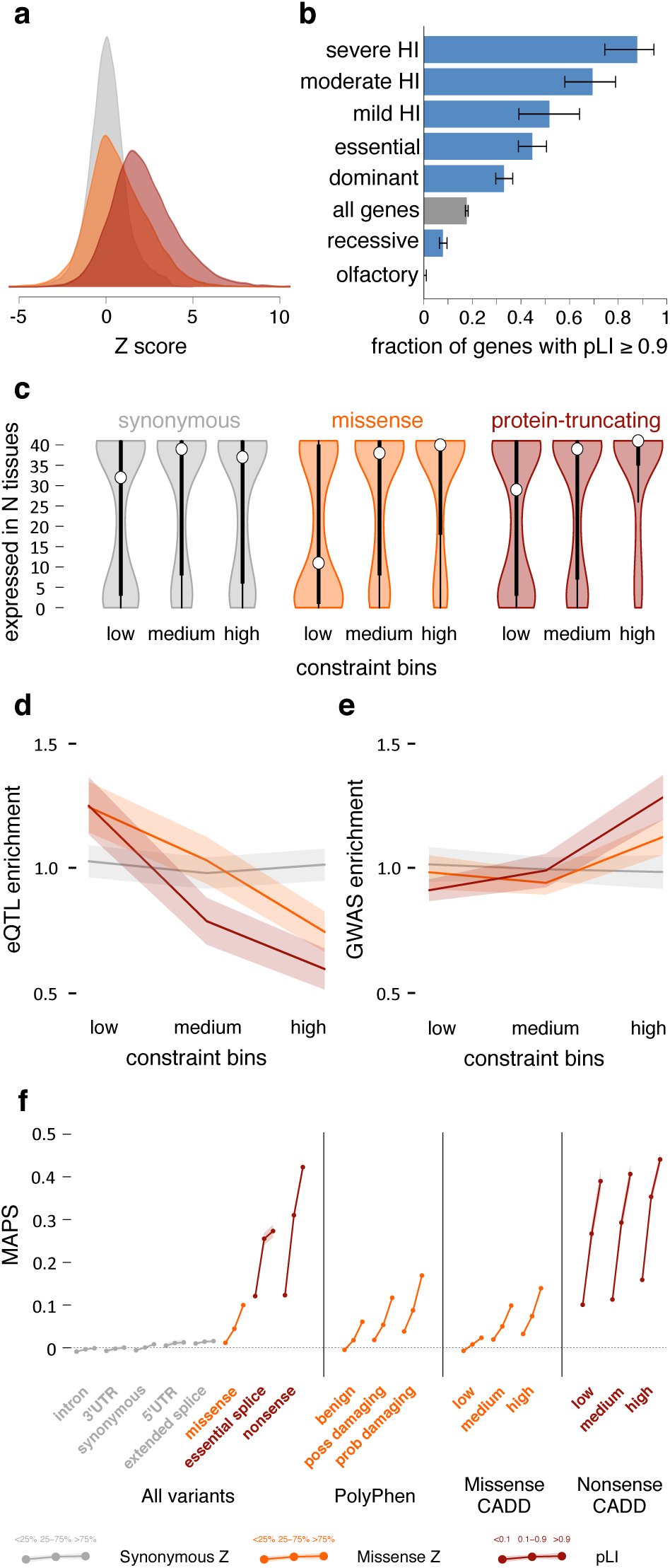
Quantifying intolerance to functional variation in genes and gene sets. a) Histograms of constraint Z scores [Samocha 2014] for 18,225 genes. This measure of departure of number of variants from expectation is normally distributed for synonymous variants, but right-shifted (higher constraint) for missense and protein-truncating variants (PTVs), indicating that more genes are intolerant to these classes of variation. b) The proportion of genes that are very likely intolerant of loss-of-function variation (pLI ≥ 0.9) is highest for ClinGen haploinsufficient genes, and stratifies by the severity and age of onset of the haploinsufficient phenotype. Genes essential in cell culture and dominant disease genes are likewise enriched for intolerant genes, while recessive disease genes and olfactory receptors have fewer intolerant genes. Black error bars indicate 95% confidence intervals (CI). c) Synonymous Z scores show no correlation with the number of tissues in which a gene is expressed, but the least missense- and PTV-constrained genes tend to be expressed in fewer tissues. Thick black bars indicate the first to third quartiles, with the white circle marking the median. d) Highly missense- and PTV-constrained genes are less likely to have eQTLs discovered in GTEx as the average gene. Shaded regions around the lines indicate 95% CI. e) Highly missense- and PTV-constrained genes are more likely to be adjacent to GWAS signals than the average gene. Shaded regions around the lines indicate 95% CI. f) MAPS (Figure 2d) is shown for each functional category, broken down by constraint score bins as shown. Missense and PTV constraint score bins provide information about natural selection at least partially orthogonal to MAPS, PolyPhen, and CADD scores, indicating that this metric should be useful in identifying variants associated with deleterious phenotypes. Shaded regions around the lines indicate 95% CI. For panels a,c-f: synonymous shown in gray, missense in orange, and protein-truncating in maroon.

Critically, we note that LoF-intolerant genes include virtually all known severe haploinsufficient human disease genes (Figure 3b), but that 72% of LoF-intolerant genes have not yet been assigned a human disease phenotype despite clear evidence for extreme selective constraint (Supplementary Information Table 13). We note that this extreme constraint does not necessarily reflect a lethal disease, but is likely to point to genes where heterozygous loss of function confers some non-trivial survival or reproductive disadvantage.

The most highly constrained missense (top 25% missense Z scores) and PTV (pLI ≥0.9) genes show higher expression levels and broader tissue expression than the least constrained genes^19^ (Figure 3c). These most highly constrained genes are also depleted for eQTLs (p < 10^−9^ for missense and PTV; Figure 3d), yet are enriched within genome-wide significant trait-associated loci (χ^2^ p < 10^−14^, Figure 3e). Intuitively, genes intolerant of PTV variation are dosage sensitive: natural selection does not tolerate a 50% deficit in expression due to the loss of single allele. Unsurprisingly, these genes are also depleted of common genetic variants that have a large enough effect on expression to be detected as eQTLs with current limited sample sizes. However, smaller changes in the expression of these genes, through weaker eQTLs or functional variants, are more likely to contribute to medically relevant phenotypes.

Finally, we investigated how these constraint metrics would stratify mutational classes according to their frequency spectrum, corrected for mutability as in the previous section (Figure 3f). The effect was most dramatic when considering nonsense variants in the LoF-intolerant set of genes. For missense variants, the missense Z score offers information additional to Polyphen2 and CADD classifications, indicating that gene-level measures of constraint offer additional information to variant-level metrics in assessing potential pathogenicity.

## ExAC improves variant interpretation in Mendelian disease

We assessed the value of ExAC as a reference dataset for clinical sequencing approaches, which typically prioritize or filter potentially deleterious variants based on functional consequence and allele frequency (AF)^6^. Filtering on ExAC reduced the number of candidate protein-altering variants by 7-fold compared to ESP, and was most powerful when the highest AF in any one population (“popmax”) was used rather than average (“global”) AF (Figure 4a). ESP is not well-powered to filter at 0.1% AF without removing many genuinely rare variants, as AF estimates based on low allele counts are both upward-biased and imprecise (Figure 4b). We thus expect that ExAC will provide a very substantial boost in the power and accuracy of variant filtering in Mendelian disease projects.

**Figure 4.**
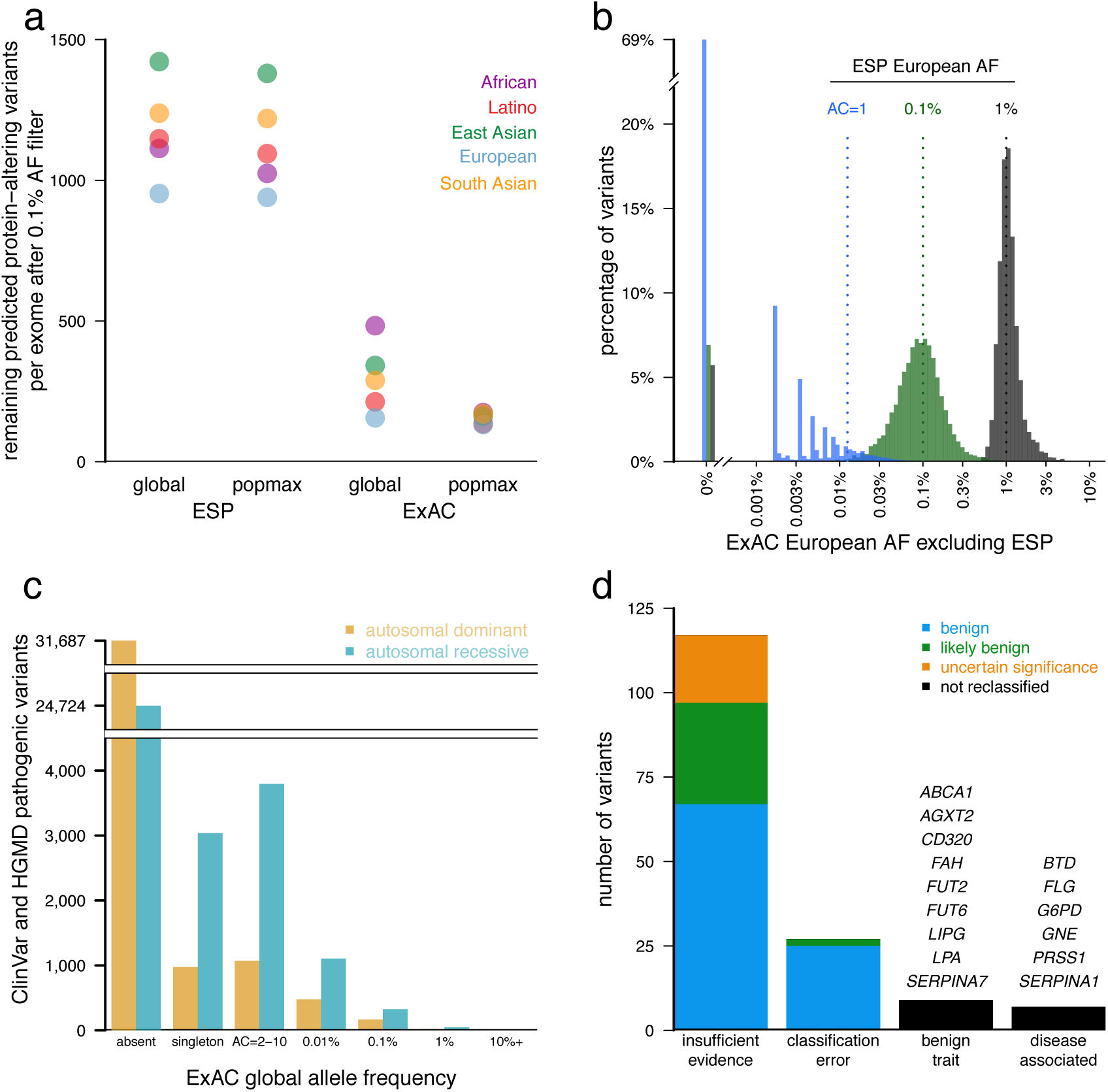
Filtering for Mendelian variant discovery. a) Predicted missense and protein-truncating variants in 500 randomly chosen ExAC individuals were filtered based on allele frequency information from ESP, or from the remaining ExAC individuals. At a 0.1% allele frequency (AF) filter, ExAC provides greater power to remove candidate variants, leaving an average of 154 variants for analysis, compared to 1090 after filtering against ESP. Popmax AF also provides greater power than global AF, particularly when populations are unequally sampled. b) Estimates of allele frequency in Europeans based on ESP are more precise at higher allele frequencies. Sampling variance and ascertainment bias make AF estimates unreliable, posing problems for Mendelian variant filtration. 69% of ESP European singletons are not seen a second time in ExAC (tall bar at left), illustrating the dangers of filtering on very low allele counts. c) Allele frequency spectrum of disease-causing variants in the Human Gene Mutation Database (HGMD) and/or pathogenic or likely pathogenic variants in ClinVar for well characterized autosomal dominant and autosomal recessive disease genes^27^. Most are not found in ExAC; however, many of the reportedly pathogenic variants found in ExAC are at too high a frequency to be consistent with disease prevalence and penetrance. d) Literature review of variants with >1% global allele frequency or >1% Latin American or South Asian population allele frequency confirmed there is insufficient evidence for pathogenicity for the majority of these variants. Variants were reclassified by ACMG guidelines^23^.

Previous large-scale sequencing studies have repeatedly shown that some purported Mendelian disease-causing genetic variants are implausibly common in the population^20–22^ (Figure 4c). The average ExAC participant harbors ~54 variants reported as disease-causing in two widely-used databases of disease-causing variants (Supplementary Information Section 5.2). Most (~41) of these are high-quality genotypes but with implausibly high (>1%) popmax AF. We therefore hypothesized that most of the supposed burden of Mendelian disease alleles per person is due not to genotyping error, but rather to misclassification in the literature and/or in databases.

We manually curated the evidence of pathogenicity for 192 previously reported pathogenic variants with AF >1% either globally or in South Asian or Latino individuals, populations that are underrepresented in previous reference databases. Nine variants had sufficient data to support disease association, typically with either mild or incompletely penetrant disease effects; the remainder either had insufficient evidence for pathogenicity, no claim of pathogenicity, or were benign traits (Supplementary Information Section 5.3). It is difficult to prove the absence of any disease association, and incomplete penetrance or genetic modifiers may contribute in some cases. Nonetheless, the high cumulative AF of these variants combined with their limited original evidence for pathogenicity suggest little contribution to disease, and 163 variants met American College of Medical Genetics criteria^23^ for reclassification as benign or likely benign (Figure 4d). 126 of these 163 have been reclassified in source databases as of December 2015 (Supplementary Information Table 20). Supporting functional data were reported for 18 of these variants, highlighting the need to review cautiously even variants with experimental support.

We also sought phenotypic data for a subset of ExAC participants homozygous for reported severe recessive disease variants, again enabling reclassification of some variants as benign. North American Indian Childhood Cirrhosis is a recessive disease of cirrhotic liver failure during childhood requiring liver transplant for survival to adulthood, previously reported to be caused by *CIRH1A* p.R565W^24^. ExAC contains 222 heterozygous and 4 homozygous Latino individuals, with a population AF of 1.92%. The 4 homozygotes had no history of liver disease and recontact in two individuals revealed normal liver function (Supplementary Information Table 22). Thus, despite the rigorous linkage and Sanger sequencing efforts that led to the original report of pathogenicity, the ExAC data demonstrate that this variant is either benign or insufficient to cause disease, highlighting the importance of matched reference populations.

The above curation efforts confirm the importance of AF filtering in analysis of candidate disease variants^6,25,26^. However, literature and database errors are prevalent even at lower AFs: the average ExAC individual contains 0.89 (<1% popmax AF) reportedly Mendelian variants in well-characterized dominant disease genes^27^ and 0.21 at <0.1% popmax AF. This inflation likely results from a combination of false reports of pathogenicity and incomplete penetrance, as we have recently shown for *PRNP*^28^. The abundance of rare functional variation in many disease genes in ExAC is a reminder that such variants should not be assumed to be causal or highly penetrant without careful segregation or case-control analysis^7,23^.

## Impact of rare protein-truncating variants

We investigated the distribution of PTVs, variants predicted to disrupt protein-coding genes through the introduction of a stop codon or frameshift or the disruption of an essential splice site; such variants are expected to be enriched for complete loss of function of the impacted genes. Naturally-occurring PTVs in humans provide a model for the functional impact of gene inactivation, and have been used to identify many genes in which LoF causes severe disease^29^, as well as rare cases where LoF is protective against disease^30^.

Among the 7,404,909 HQ variants in ExAC, we found 179,774 high-confidence PTVs (as defined in Supplementary Information Section 6), 121,309 of which are singletons. This corresponds to an average of 85 heterozygous and 35 homozygous PTVs per individual (Figure 5a). The diverse nature of the cohort enables the discovery of substantial numbers of novel PTVs: out of 58,435 PTVs with an allele count greater than one, 33,625 occur in only one population. However, while PTVs as a category are extremely rare, the majority of the PTVs found in any one person are common, and each individual has only ~2 singleton PTVs, of which 0.14 are found in PTV-constrained genes (pLI >0.9). ExAC recapitulates known aspects of population demographic models, including an increase in intermediate-frequency (1-5%) PTVs in Finland^31^ and relatively common (>1%) PTVs in Africans (Figure 5b). However, these differences are diminished when considering only LoF-constrained (pLI > 0.9) genes (Extended Data Figure 10).

**Figure 5.**
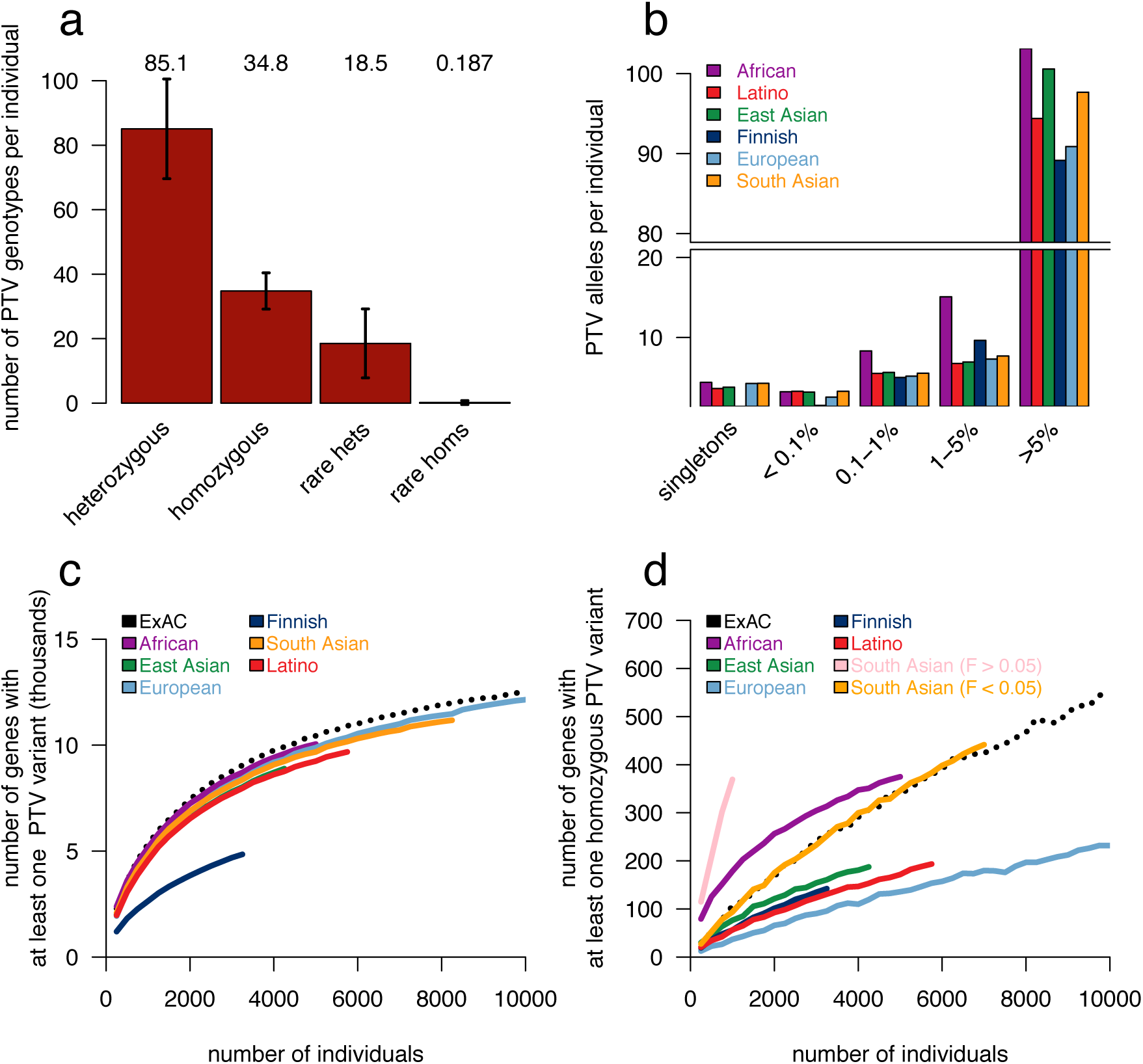
Protein-truncating variation in ExAC. a) The average ExAC individual has 85 heterozygous and 35 homozygous protein-truncating variants (PTVs), of which 18 and 0.19 are rare (<0.1% popmax AF), respectively. Error bars represent standard deviation. b) Breakdown of PTVs per individual (a) by popmax AF bin. Across all populations, most PTVs found in a given individual are common (>5% popmax AF). c-d) Number of genes with at least one PTV (c) or homozygous PTV (d) as a function of number of individuals, downsampled from ExAC. South Asian population is broken down by consanguinity (Inbreeding coefficient, F).

Using a sub-sampling approach, we show that the discovery of both heterozygous (Figure 5c) and homozygous (Figure 5d) PTVs scales very differently across human populations, with implications for the design of large-scale sequencing studies for the ascertainment of human “knockouts” described below.

## Discussion

Here we describe the generation and analysis of the most comprehensive catalogue of human protein-coding genetic variation to date, incorporating high-quality exome sequencing data from 60,706 individuals of diverse geographic ancestry. The resulting call set provides unprecedented resolution for the analysis of low-frequency protein-coding variants in human populations, as well as a public resource [exac.broadinstitute.org] for the clinical interpretation of genetic variants observed in disease patients.

The very large sample size of ExAC also provides opportunities for a high-resolution analysis of the sensitivity of human genes to functional variation. While previous sample sizes have been adequately powered for the assessment of gene-level intolerance to missense variation^11,14^, ExAC provides for the first time sufficient power to investigate genic intolerance to PTVs, highlighting 3,230 highly LoF-intolerant genes, 72% of which have no established human disease phenotype in OMIM or ClinVar. We note that this extreme constraint does not necessarily reflect a lethal disease, but is likely to point to genes where heterozygous loss of function confers some non-trivial survival or reproductive disadvantage. In independent work [Ruderfer et al., manuscript submitted] we show that ExAC similarly provides power to identify genes intolerant of copy number variation. Quantification of genic intolerance to both classes of variation will provide added power to disease studies.

The ExAC resource provides the largest database to date for the estimation of allele frequency for protein-coding genetic variants, providing a powerful filter for analysis of candidate pathogenic variants in severe Mendelian diseases. Frequency data from ESP^1^ have been widely used for this purpose, but those data are limited by population diversity and by resolution at allele frequencies ≤0.1%. ExAC therefore provides substantially improved power for Mendelian analyses, although it is still limited in power at lower allele frequencies, emphasizing the need for more sophisticated pathogenic variant filtering strategies alongside on-going data aggregation efforts.

Finally, we show that different populations confer different advantages in the discovery of gene-disrupting PTVs, providing guidance for the identification of human “knockouts” to understand gene function. Sampling multiple populations would likely be a fruitful strategy for a researcher investigating common PTV variation. However, discovery of homozygous PTVs is markedly enhanced in the South Asian samples, which come primarily from a Pakistani cohort with 38.3% of individuals self-reporting as having closely related parents, emphasizing the extreme value of consanguineous cohorts for “human knockout” discovery^32–34^ (Figure 5d). Other approaches to enriching for homozygosity of rare PTVs, such as focusing on bottlenecked populations, have already proved fruitful^31,32^.

Even with this large collection of jointly processed exomes, many limitations remain. Firstly, most ExAC individuals were ascertained for biomedically important disease; while we have attempted to exclude severe pediatric diseases, the inclusion of both cases and controls for several polygenic disorders means that ExAC certainly contains disease-associated variants^35^. Secondly, future reference databases would benefit from including a broader sampling of human diversity, especially from underrepresented Middle Eastern and African populations. Thirdly, the inclusion of whole genomes will also be critical to investigate additional classes of functional variation and identify non-coding constrained regions. Finally, and most critically, detailed phenotype data are unavailable for the vast majority of ExAC samples; future initiatives that assemble sequence and clinical data from very large-scale cohorts will be required to fully translate human genetic findings into biological and clinical understanding.

While the ExAC dataset exceeds the scale of previously available frequency reference datasets, much remains to be gained by further increases in sample size. Indeed, the fact that even the rarest transversions have mutational rates^11^ on the order of 1 × 10^−9^ implies that the vast majority of possible non-lethal SNVs likely exist in some living human. ExAC already includes >63% of all possible protein-coding CpG transitions at well-covered synonymous sites; orders-of-magnitude increases in sample size will eventually lead to saturation of other classes of variation.

ExAC was made possible by the willingness of multiple large disease-focused consortia to share their raw data, and by the availability of the software and computational resources required to create a harmonized variant call set on the scale of tens of thousands of samples. The creation of yet larger reference variant databases will require continued emphasis on the value of genomic data sharing.

## Online Methods

### Variant discovery

We assembled approximately 1 petabyte of raw sequencing data (FASTQ files) from 91,796 individual exomes drawn from a wide range of primarily disease-focused consortia (Supplementary Information Table 2). We processed these exomes through a single informatic pipeline and performed joint variant calling of single nucleotide variants (SNVs) and short insertions and deletions (indels) across all samples using a new version of the Genome Analysis Toolkit (GATK) HaplotypeCaller pipeline. Variant discovery was performed within a defined exome region that includes Gencode v19 coding regions and flanking 50 bases. At each site, sequence information from all individuals was used to assess the evidence for the presence of a variant in each individual. Full details of data processing, variant calling and resources are described in the Supplementary Information Sections 1.1-1.4.

### Quality assessment

We leveraged a variety of sources of internal and external validation data to calibrate filters and evaluate the quality of filtered variants (Supplementary Information Table 7). We adjusted the standard GATK variant site filtering^36^ to increase the number of singleton variants that pass this filter, while maintaining a singleton transmission rate of 50.1%, very near the expected 50%, within sequenced trios. We then used the remaining passing variants to assess depth and genotype quality filters compared to >10,000 samples that had been directly genotyped using SNP arrays (Illumina HumanExome) and achieved 97-99% heterozygous concordance, consistent with known error rates for rare variants in chip-based genotyping^37^. Relative to a “platinum standard” genome sequenced using five different technologies^38^, we achieved sensitivity of 99.8% and false discovery rates (FDR) of 0.056% for single nucleotide variants (SNVs), and corresponding rates of 95.1% and 2.17% for insertions and deletions (indels). Lastly, we compared 13 representative Non-Finnish European exomes included in the call set with their corresponding 30x PCR-Free genome. The overall SNV and indel FDR was 0.14% and 4.71%, while for SNV singletons was 0.389%. The overall FDR by annotation classes missense, synonymous and protein truncating variants (including indels) were 0.076%, 0.055% and 0.471% respectively (Supplementary Information Table 5 and 6). Full details of quality assessments are described in the Supplementary Information Section 1.6.

### Sample filtering

The 91,796 samples were filtered based on two criteria. First, samples that were outliers for key metrics were removed (Extended Data Figure 2b). Second, in order to generate allele frequencies based on independent observations without enrichment of Mendelian disease alleles, we restricted the final release data set to unrelated adults with high-quality sequence data and without severe pediatric disease. After filtering, only 60,706 samples remained, consisting of ~77% of Agilent (33 Mb target) and ~12% of Illumina (37.7 Mb target) exome captures. Full details of the filtering process are described in the Supplementary Information Section 1.7.

### ExAC data release

For each variant, summary data for genotype quality, allele depth and population specific allele counts were calculated before removing all genotype data. This variant summary file was then functionally annotated using variant effect predictor (VEP) with the LOFTEE plugin. This data set can be accessed via the ExAC Browser (http://exac.broadinstitute.org) or downloaded from ftp://ftp.broadinstitute.org/pub/ExAC_release/release0.3/ExAC.r0.3.sites.vep.vcf.gz. Full details regarding the annotation of the ExAC data set are described in the Supplementary Information Sections 1.9-1.10.

## Acknowledgements

We would like to thank the reviewers and editor for their time, valuable comments and suggestions. The scientific community for their support and comments on biorxiv, twitter and other public forums. Brendan Bulik-Sullivan and Jon Bloom for their help with mathematical notation.

M.Lek is supported by the Australian National Health and Medical Research Council CJ Martin Fellowship, Australian American Association Sir Keith Murdoch Fellowship and the MDA/AANEM Development Grant. K.J.K. is supported by NIGMS Fellowship (F32GM115208). A.H.O. is supported by Pfizer/ACMG Foundation Translational Genomic Fellowship. J.S.W. is supported by Fondation Leducq and Wellcome Trust. A.J.H. is supported by NSF Graduate Research Fellowship. M.I.K is supported by Instrumentarium Science Foundation, Finland; Finnish Foundation for Cardiovascular Research; Orion Research Foundation and the University of Eastern Finland, Saastamoinen Foundation. P.N. is supported by John S. LaDue Memorial Fellowship in Cardiology, Harvard Medical School. G.M.P. is supported by the National Heart, Lung, and Blood Institute of the National Institutes of Health under Award Number K01HL125751. M.T.Tusie-Luna is supported by CONACyT grant 128877. H.W. is supported by postdoctoral award from the American Heart Association (15POST23280019). R.E. is supported by Instituto Salud Carlos III-FIS-FEDER-ERDF: RD12/0042/0013, PI12/00232; Agència de Gestió Ajuts Universitaris de Recerca: 2014 SGR 240. S.K. is supported by grants from the National Institutes of Health (R01HL107816), the Donovan Family Foundation and Fondation Leducq. S.J.G. is supported by NIH/NIMH grant R01MH085521 and NARSAD: The Brain and Behavior Research Foundation and the Sidney R. Baer, Jr. Foundation. M.I.M is supported by Wellcome Trust Senior Investigator, NIHR Senior Investigator;; EU Framework VII HEALTH-F4-2007-201413; Medical Research Council G0601261; Wellcome Trust 090532, 098381, 090367; NIH RC2-DK088389, U01-DK085545. R.M is supported by Canadian Institutes of Health Research MOP136936; MOP82810, MOP77682, Canadian Foundation for Innovation 11966, Heart & Stroke Foundation of Canada T-7268. J.M.S is supported by NINDS grants NS40024-09S1 and NS085048. P.S. is supported by NIMH grant MH095034 and MH089905. P.F.S is supported by Swedish Research Council award D0886501; NIMH grants MH077139 and MH094421; Yeargen Family; Stanley Center. H.C.W. is supported by BHF Centre of Research Excellence, NIHR Senior Investigator. M.T.Tsuang is supported by NIH/NIMH grant R01MH085560. D.G.M is supported by NIGMS R01 GM104371 and NIDDK U54 DK105566.

**ATVB & Precocious Coronary Artery Disease Study (PROCARDIS)**: Exome sequencing was supported by a grant from the NHGRI (5U54HG003067-11) to Drs. Gabriel and Lander. **Bulgarian Trios**: Medical Research Council (MRC) Centre (G0800509) and ProgramGrants (G0801418), the European Community’s Seventh Framework Programme (HEALTH-F2-2010-241909 (Project EU-GEI)),and NIMH(2P50MH066392-05A1). **GoT2D & T2DGENES**: NHGRI (“Large Scale Sequencing and Analysis of Genomes” U54HG003067), NIDDK (“Multiethnic Study of Type 2 Diabetes Genes” U01DK085526), NIH (“LowF Pass Sequencing and High Density SNP Genotyping in Type 2 Diabetes” 1RC2DK088389), National Institutes of Health (“Multiethnic Study of Type 2 Diabetes Genes” U01s DK085526, DK085501, DK085524, DK085545, DK085584; “LowF Pass Sequencing and HighF Density SNP Genotyping for Type 2 Diabetes” DK088389). The German Center for Diabetes Research (DZD). National Institutes of Health (RC2F DK088389, DK085545, DK098032). Wellcome Trust (090532, 098381). National Institutes of Health (R01DK062370, R01DK098032, RC2DK088389). **METSIM**: Academy of Finland and the Finnish Cardiovascular Research Foundation. **Inflammatory Bowel Disease**: The Helmsley Trust Foundation, #2015PG-IBD001, Large Scale Sequencing and Analysis of Genomes Grant (NHGRI), 5 U54 HG003067-13. Jackson Heart Study: We thank the **Jackson Heart Study** (JHS) participants and staff for their contributions to this work. The JHS is supported by contracts HHSN268201300046C, HHSN268201300047C, HHSN268201300048C, HHSN268201300049C, HHSN268201300050C from the National Heart, Lung, and Blood Institute and the National Institute on Minority Health and Health Disparities. **Ottawa Genomics Heart Study**: Canadian Institutes of Health Research MOP136936; M0P82810, MOP77682, Canadian Foundation for Innovation 11966, Heart & Stroke Foundation of Canada T-7268. Exome sequencing was supported by a grant from the NHGRI (5U54HG003067-11) to Drs. Gabriel and Lander.

**Pakistan Risk of Myocardial Infarction Study (PROMIS)**: Exome sequencing was supported by a grant from the NHGRI (5U54HG003067-11) to Drs. Gabriel and Lander. Fieldwork in the study has been supported through funds available to investigators at the Center for Non-Communicable Diseases, Pakistan and the University of Cambridge, UK.

**Registre Gironi del COR (REGICOR)**: Spanish Ministry of Economy and Innovation through the Carlos III Health Institute [Red HERACLES RD12/0042, CIBER Epidemiología y Salud Pública, PI12/00232, PI09/90506, PI08/1327, PI05/1251, PI05/1297], European Funds for Development (ERDF-FEDER), and by the Catalan Research and Technology Innovation Interdepartmental Commission [SGR 1195].

**Swedish Schizophrenia & Bipolar Studies**: National Institutes of Health (NIH)/National Institute of Mental Health (NIMH) ARRA Grand Opportunity grant NIMHRC2MH089905, the Sylvan Herman Foundation, the Stanley Center for Psychiatric Research, the Stanley Medical Research Institute, NIH/National Human GenomeResearch Institute (NHGRI) grant U54HG003067. **SIGMA-T2D**: The work was conducted as part of the Slim Initiative for Genomic Medicine, a project funded by the Carlos Slim Health Institute in Mexico. The UNAM/INCMNSZ Diabetes Study was supported by Consejo Nacional de Ciencia y Tecnologiía grants 138826, 128877, CONACT-SALUD 2009-01-115250, and a grant from Dirección General de Asuntos del Personal Académico, UNAM, IT 214711. The Diabetes in Mexico Study was supported by Consejo Nacional de Ciencia y Tecnología grant 86867 and by Instituto Carlos Slim de la Salud, A.C. The Mexico City Diabetes Study was supported by National Institutes of Health (NIH) grant R01HL24799 and by the Consejo Nacional de Ciencia y Tenologia grants 2092, M9303, F677-M9407, 251M, and 2005-C01-14502, SALUD 2010-2-151165. **Schizophrenia Trios from Taiwan**: NIH/NIMH grant R01MH085560. **Tourette Syndrome Association International Consortium for Genomics (TSAICG)**: NIH/NINDS U01 NS40024-09S1. **Exome Aggregation Consoritum (ExAC)**: NIDDK U54 DK105566.

## Author Contributions

M.Lek,K.J.K.,E.V.M.,K.E.S.,E.B.,T.F.,A.H.O.,J.S.W.,A.J.H.,B.B.C.,T.T.,D.P.B.,J.A.K.,L.D.,K.E.,F.Z.,J.Z.,E.P.,M.J.D.,D.G.M. contributed to the analysis and writing of the manuscript. M.Lek, E.B.,T.F.,K.J.K.,E.V.M.,F.Z.,D.P.B.,J.B.,D.N.C.,N.D.,M.D.,R.D.,J.F.,M.F.,L.G.,J.G.,N.G.,D.H.,A.K.,M.I.K.,A.L.M.,P.N.,L.O.,G.M.P.,R.P.,M.A.R.,V.R.,S.A.R.,D.M.R.,K.S.,P.D.S.,C.S.,B.P.T.,G.T.,M.T.T.,B.W.,H.W.,D.Y.,S.B.G.,M.J.D.,D.G.M.contributed to the production of the ExAC data set. D.M.A.,D.A.,M.B.,J.D.,S.D.,R.E.,J.C.F.,S.B.G.,G.G.,S.J.G.,C.M.H.,S.K.,M.Laakso,S.M.,M.I.M.,D.M.,R.M.,B.M.N.,A.P.,S.M.P.,D.S.,J.S.,P.S.,P.F.S.,J.T.,M.T.T.,H.C.W.,J.G.W.,M.J.D.,D.G.M. contributed to the design and conduct of the various exome sequencing studies and critical review of manuscript.

## Author Information

P.F.S is a scientific advisor to Pfizer.

ExAC data set is publicly available at http://exac.broadinstitute.org

## Collaborators (alphabetical order)

Hanna E Abboud^61^, Goncalo Abecasis^35^, Carlos A Aguilar-Salinas^62^, Olimpia Arellano-Campos^62^, Gil Atzmon^63,64^, Ingvild Aukrust^65,66,67^, Cathy L Barr^68,69^, Graeme I Bell^70^, Graeme I Bell^70,71^, Sarah Bergen^42^, Lise Bjørkhaug^66,67^, John Blangero^72,73^, Donald W Bowden^74,75,76^, Cathy L Budman^77^, Noël P Burtt^2^, Federico Centeno-Cruz^78^, John C Chambers^79,80,81^, Kimberly Chambert^6^, Robert Clarke^82^, Rory Collins^82^, Giovanni Coppola^83^, Emilio J Córdova^78^, Maria L Cortes^18^, Nancy J Cox^84^, Ravindranath Duggirala^85^, Martin Farrall^59,44^, Juan C Fernandez-Lopez^78^, Pierre Fontanillas^2^, Timothy M Frayling^86^, Nelson B Freimer^83^, Christian Fuchsberger^35^, Humberto García-Ortiz^78^, Anuj Goel^59,44^, María J Gómez-Vázquez^62^, María E González-Villalpando^87^, Clicerio González-Villalpando^87^, Marco A Grados^88^, Leif Groop^89^, Christopher A Haiman^90^, Craig L Hanis^91^, Craig L Hanis^91^, Andrew T Hattersley^86^, Brian E Henderson^92^, Jemma C Hopewell^82^, Alicia Huerta-Chagoya^93^, Sergio Islas-Andrade^94^, Suzanne BR Jacobs^2^, Shapour Jalilzadeh^59,44^, Christopher P Jenkinson^61^, Jennifer Moran^2^, Silvia Jiménez-Morale^78^, Anna Kähler^42^, Robert A King^95^, George Kirov^96^, Jaspal S Kooner^80,9,81^, Theodosios Kyriakou^59,44^, Jong-Young Lee^97^, Donna M Lehman^61^, Gholson Lyon^98^, William MacMahon^99^, Patrik KE Magnusson^42^, Anubha Mahajan^100^, Jaume Marrugat^37^, Angélica Martínez-Hernández^78^, Carol A Mathews^101^, Gilean McVean^100^, James B Meigs^102,26^, Thomas Meitinger^103,104^, Elvia Mendoza-Caamal^78^, Josep M Mercader^2,105,106^, Karen L Mohlke^55^, Hortensia Moreno-Macías^107^, Andrew P Morris^108,100,109^, Laeya A Najmi^65,110^, Pal R Nj0lstad^65,66^, Michael C O'Donovan^96^, Maria L Ordóñez-Sánchez^62^, Michael J Owen^96^, Taesung Park^111,112^, David L Pauls^25^, Danielle Posthuma^113,114,115^, Cristina Revilla-Monsalve^94^, Laura Riba^93^, Stephan Ripke^6^, Rosario Rodríguez-Guillén^62^, Maribel Rodríguez-Torres^62^, Paul Sandor^116,68^, Mark Seielstad^117,118^, Rob Sladek^119,120,121^, Xavier Soberón^78^, Timothy D Spector^122^, Shyong E Tai^123,124,125^, Tanya M Teslovich^35^, Geoffrey Walford^105,26^, Lynne R Wilkens^92^, Amy L Williams^2,126^

## Footnotes

61 Department of Medicine, University of Texas Health Science Center, San Antonio, TX, USA

62 Instituto Nacional de Ciencias M_dicas y Nutrici—n Salvador Zubir‡n, Mexico City, Mexico

63 Departments of Medicine and Genetics, Albert Einstein College of Medicine, New York City, NY, USA

64 Department of Natural Science, University of Haifa, Haifa, Israel

65 Department of Clinical Science, University of Bergen, Bergen, Norway

66 Department of Pediatrics, Haukeland University Hospital, Bergen, Norway

67 Department of Biomedicine, University of Bergen, Bergen, Norway

68 The Toronto Western Research Institute, University Health Network, Toronto, Canada

69 The Hospital for Sick Children, Toronto, Canada

70 Departments of Medicine and Human Genetics, University of Chicago, Chicago, IL, USA

71 Department of Medicine, University of Chicago, Chicago, IL, USA

72 South Texas Diabetes and Obesity Institute, University of Texas Health Science Center, San Antonio, TX, USA

73 University of Texas Rio Grande Valley, Brownsville, TX, USA

74 Department of Biochemistry, Wake Forest School of Medicine, Winston-Salem, NC, USA

75 Center for Genomics and Personalized Medicine Research, Wake Forest School of Medicine, Winston-Salem, NC, USA

76 Center for Diabetes Research, Wake Forest School of Medicine, Winston-Salem, NC, USA

77 North Shore-Long Island Jewish Health System, Manhasset, NY, USA

78 Instituto Nacional de Medicina Gen—mica, Mexico City, Mexico

79 Department of Epidemiology and Biostatistics, Imperial College London, London, UK

80 Department of Cardiology, Ealing Hospital NHS Trust, Southall, UK

81 Imperial College Healthcare NHS Trust, Imperial College London, London, UK

82 Nuffield Department of Population Health, University of Oxford, Oxford, UK

83 Center for Neurobehavioral Genetics, University of California, Los Angeles, CA, USA

84 Vanderbilt Genetics Institute, Vanderbilt University School of Medicine, Nashville, TN, USA

85 Department of Genetics, Texas Biomedical Research Institute, San Antonio, TX, USA

86 University of Exeter Medical School, University of Exeter, Exeter, UK

87 Instituto Nacional de Salud Publica, Mexico City, Mexico

88 Department of Psychiatry and Behavioral Sciences, Johns Hopkins University School of Medicine, Baltimore, MD, USA

89 Department of Clinical Sciences, Lund University Diabetes Centre, Malm_, Sweden

90 Department of Preventive Medicine, University of Southern California, Los Angeles, CA, USA

91 Human Genetics Center, The University of Texas Health Science Center, Houston, TX, USA

92 Epidemiology Program, University of Hawaii Cancer Center, Honolulu, HI, USA

93 Instituto de Investigaciones Biom_dicas, Mexico City, Mexico

94 Instituto Mexicano del Seguro Social, Mexico City, Mexico

95 Department of Genetics, Yale University School of Medicine, New Haven, CT, USA

96 MRC Centre for Neuropsychiatric Genetics and Genomics, Cardiff University, Cardiff, UK

97 Center for Genome Science, Korea National Institute of Health, Chungcheongbuk-do, Republic of Korea

98 Stanley Institute for Cognitive Genomics, Cold Spring Harbor Laboratory, Woodbury, NY, USA

99 Department of Psychiatry, University of Utah, Salt Lake City, UT, USA

100 Nuffield Department of Medicine, University of Oxford, Oxford, UK

101 Department of Psychiatry, University of Florida, Gainesville, FL, USA

102 General Medicine Division, Massachusetts General Hospital, Boston, MA, USA

103 Institute of Human Genetics, Technische Universit_t MŸnchen, Munich, Germany

104 Institute of Human Genetics, German Research Center for Environmental Health, Neuherberg, Germany

105 Diabetes Research Center (Diabetes Unit), Massachusetts General Hospital, Boston, MA, USA

106 Research Program in Computational Biology, Barcelona Supercomputing Center, Barcelona, Spain

107 Universidad Aut—noma Metropolitana, Mexico City, Mexico

108 Estonian Genome Centre,University of Tartu,Tartu,Estonia, University of Tartu, Tartu, Estonia

109 Department of Biostatistics, University of Liverpool, Liverpool, UK

110 Center for Medical Genetics and Molecular Medicine, Haukeland University Hospital, Bergen, Norway

111 Interdisoiplinary Program in Bioinformatics, Seoul National University, Seoul, Republic of Korea

112 Department of Statistics, Seoul National University, Seoul, Republic of Korea

113 Department of Functional Genomics, University of Amsterdam, Amsterdam, The Netherlands

114 Department of Clinical Genetics, VU Medical Centre, Amsterdam, The Netherlands

115 Department of Child and Adolescent Psychiatry, Erasmus University Medical Centre, Rotterdam, The Netherlands

116 Department of Psychiatry, University of Toronto, Toronto, Canada

117 Department of Laboratory Medicine, University of California, San Francisco, CA, USA

118 Blood Systems Research Institute, San Francisco, CA, USA

119 Department of Human Genetics, McGill University, Montreal, Canada

120 Department of Medicine, McGill University, Montreal, Canada

121 McGill University and G_nome Qu_bec Innovation Centre, Montreal, Canada

122 Department of Twin Research and Genetic Epidemiology, King's College London, London, UK

123 Saw Swee Hock School of Public Health, National University of Singapore, Singapore, Singapore

124 Department of Medicine, National University of Singapore, Singapore, Singapore

125 Cardiovascular & Metabolic Disorders Program, Duke-NUS Graduate Medical School Singapore, Singapore, Singapore

126 Department of Biological Sciences, Columbia University, New York, NY, USA

---

**Extended Data Figure 1.**
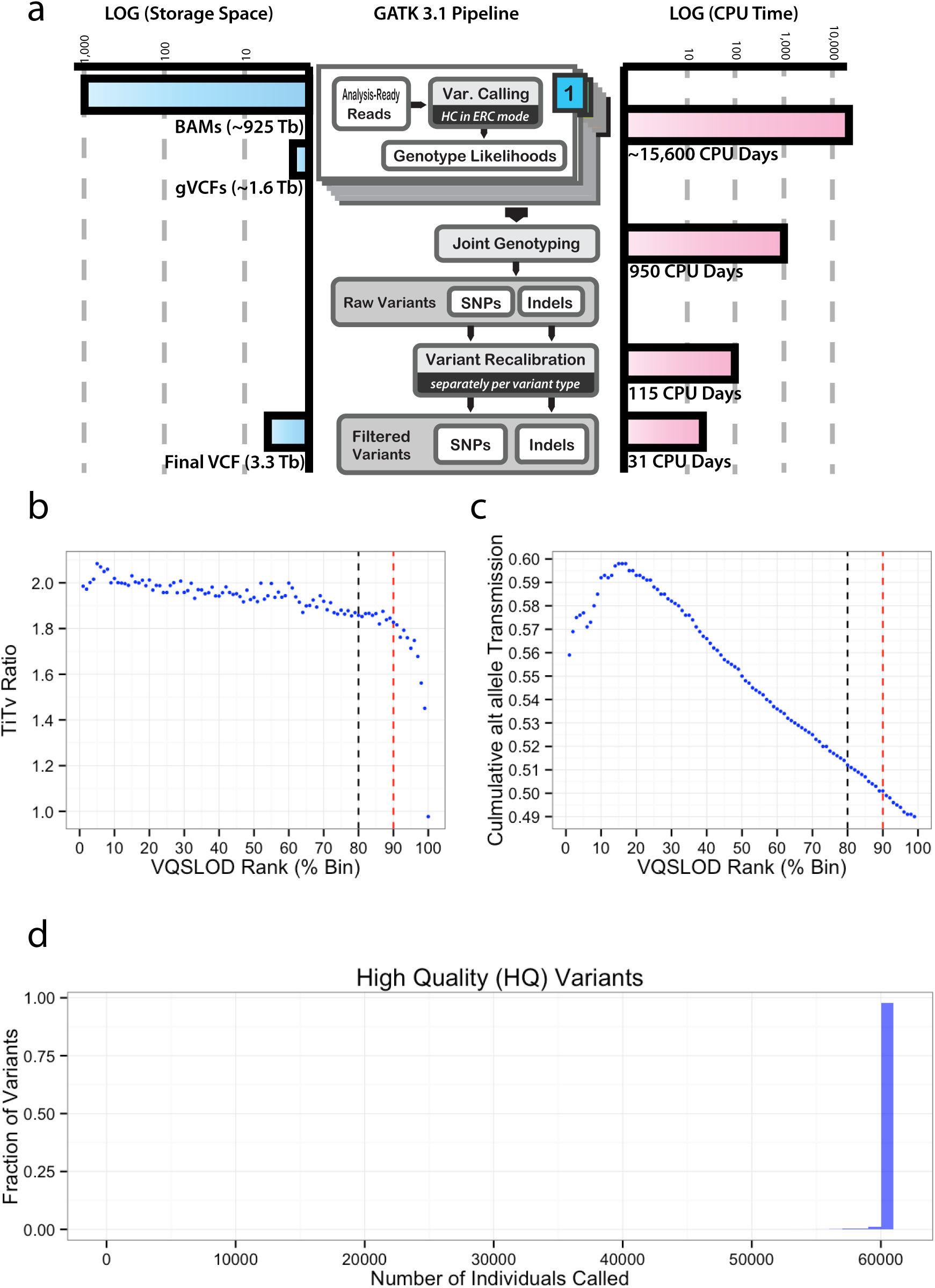
The GATK 3.1 pipeline used for the joint calling of 91,796 exomes. a) The resources used for the variant calling in terms of CPU days and storage (terabytes). b) The impact of VQSLOD on singleton TiTv. c) The impact of VQSLOD on singleton transmission in trios. Note: In b) and c) singleton variants discovered in joint called set was ordered by VQSLOD in descending order (i.e. higher confident variants first) and then binned into percentiles. The black dotted line indicates the current VQSR cut off and the red dotted line is where the less stringent threshold was moved. d) The number of individuals called at each variant site as a fraction of the total number of High Quality (HQ) variants.

**Extended Data Figure 2.**
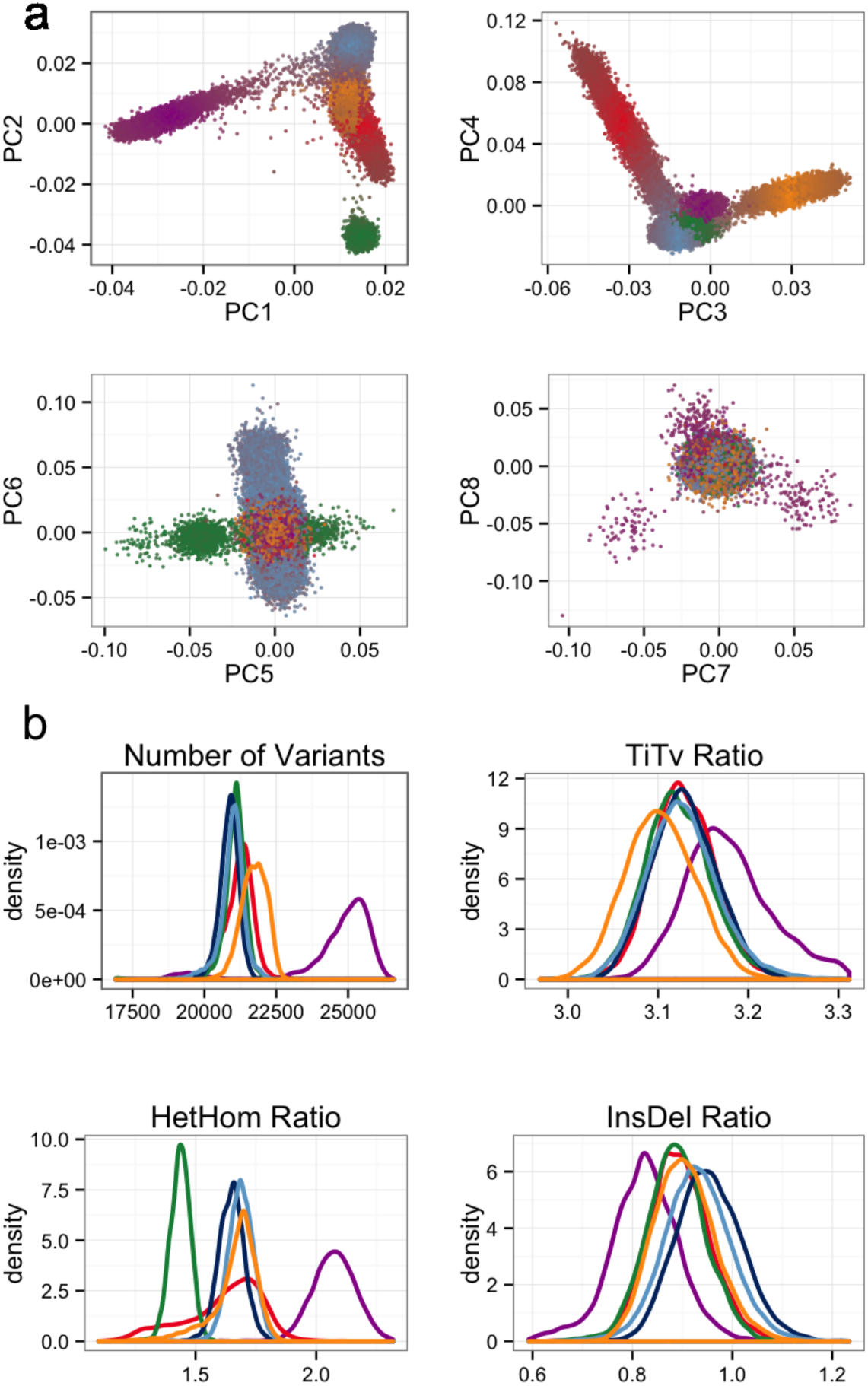
Principal component analysis (PCA) and key metrics used to filter samples. a) Principal component analysis using a set of 5,400 common exome SNPs. Individuals are colored by their distance from each of the population cluster centers using the first 4 principal components. b) The metrics number of variants, TiTv, alternate heterozygous/homozygous (HetHom) ratio and Insertion/Deletion (InsDel) ratio. Populations are their respective colors: Latino (red), African (purple), European (blue), South Asian (yellow) and East Asian (green).

**Extended Data Figure 3.**
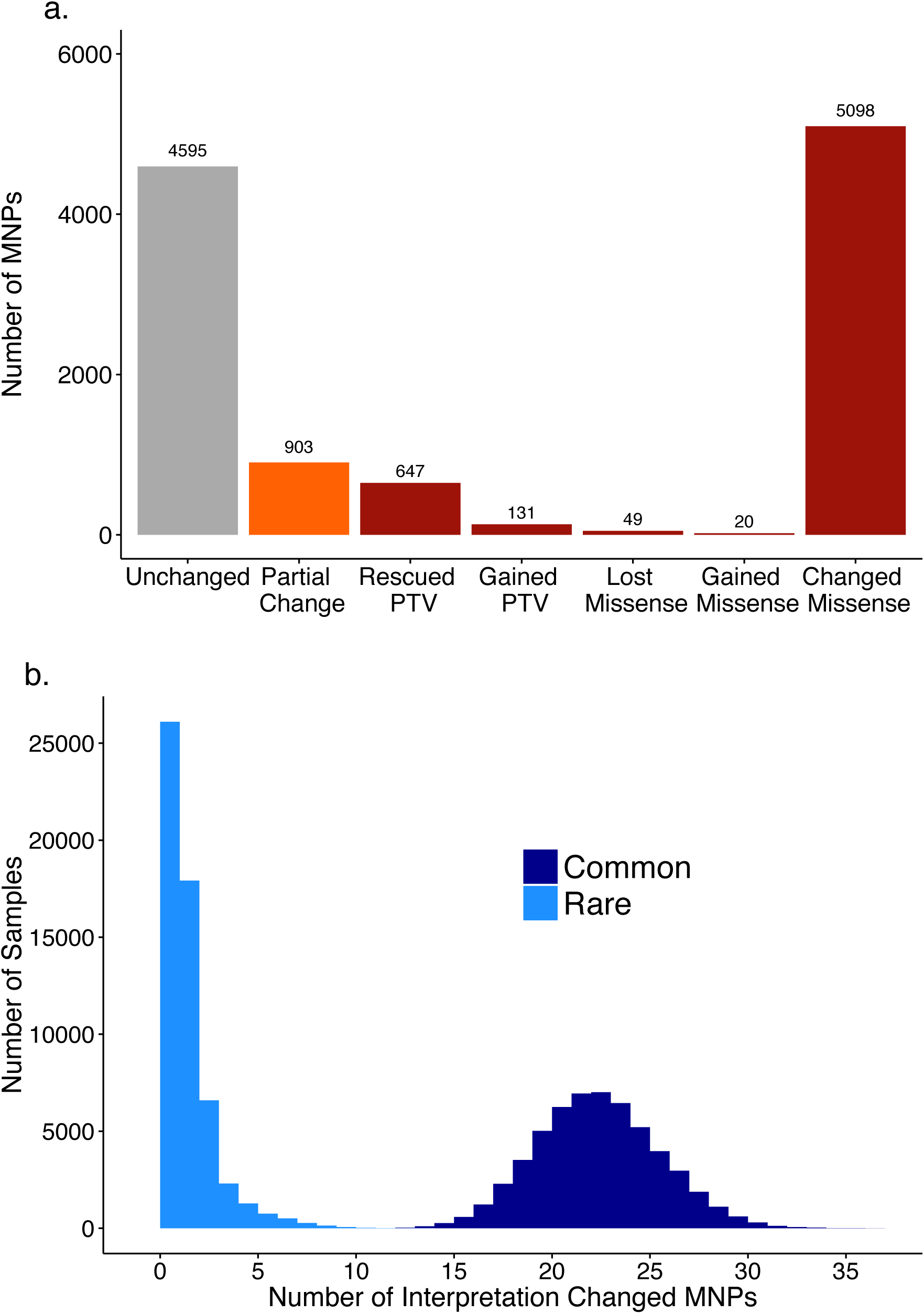
Multi-nucleotide variants discovered in the ExAC data set. a) Number of MNPs per impact on the variant interpretation. b) Distribution of the number of MNPs per sample where phasing changes interpretation, separated by allele frequency. Common > 1%, Rare < 1%. MNPs comprised of a rare and common allele are considered rare as this defines the frequency of the MNP.

**Extended Data Figure 4.**
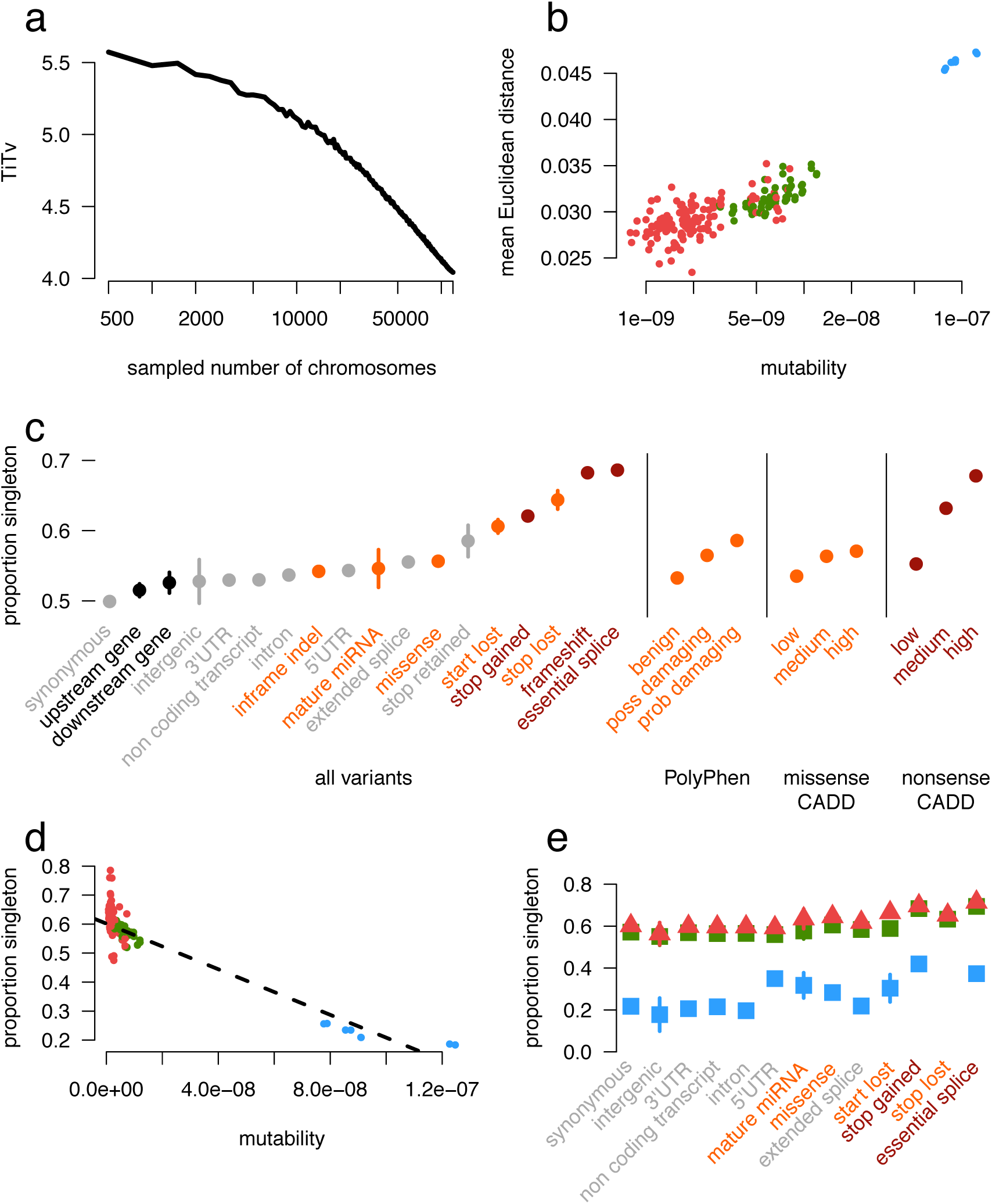
The impact of recurrence across different mutation and functional classes. a) TiTv (Transition to transversion) ratio of synonymous variants at downsampled intervals of ExAC. The TiTv is relatively stable at previous sample sizes (<5000) but changes drastically at larger sample sizes. b) For synonymous doubleton variants, mutability of each trinucleotide context is correlated with mean Euclidean distance of individuals that share the doubleton. Transversion (red) and non-CpG transition (green) doubletons are more likely to be found in closer PCA space (i.e. more similar ethnicities) than CpG transitions (blue) c) The proportion singleton among various functional categories. The functional category stop lost has a higher singleton rate than nonsense. Error bars represent standard error of the mean. d) Among synonymous variants, mutability of each trinucleotide context is correlated with proportion singleton, suggesting CpG transitions (blue) are more likely to have multiple independent origins driving their allele frequency up. e) The proportion singleton metric from c) broken down by transversions, non-CpG transitions, and CpG variants. Notably, there is a wide variation in singleton rates among mutational contexts in functional classes, and there are no stop-lost CpG transitions. Error bars represent standard error of the mean.

**Extended Data Figure 5.**
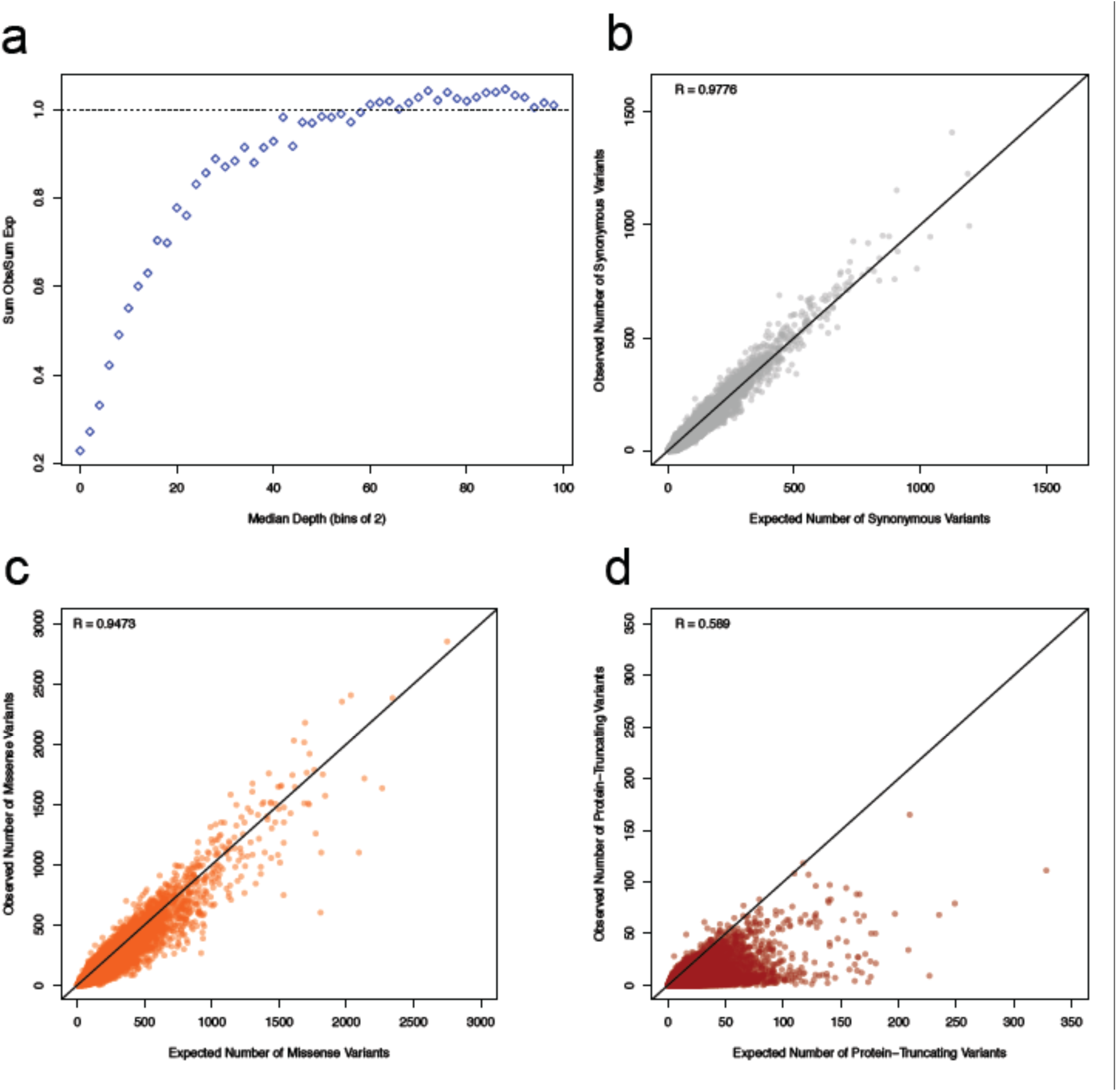
Relationships between depth and observed vs expected variants as well as correlations between observed and expected variant counts for synonymous, missense, and protein-truncating. a) The relationship between the median depth of exons (bins of 2) and the sum of all observed synonymous variants in those exons divided by the sum of all expected synonymous variants. The curve was used to determine the appropriate depth adjustment for expected variant counts. For the rest of the panels, the correlation between the depth-adjusted expected variants counts and observed are depicted for synonymous (b), missense (c), and protein-truncating (d). The black line indicates a perfect correlation (slope = 1). Axes have been trimmed to remove *TTN*.

**Extended Data Figure 6.**
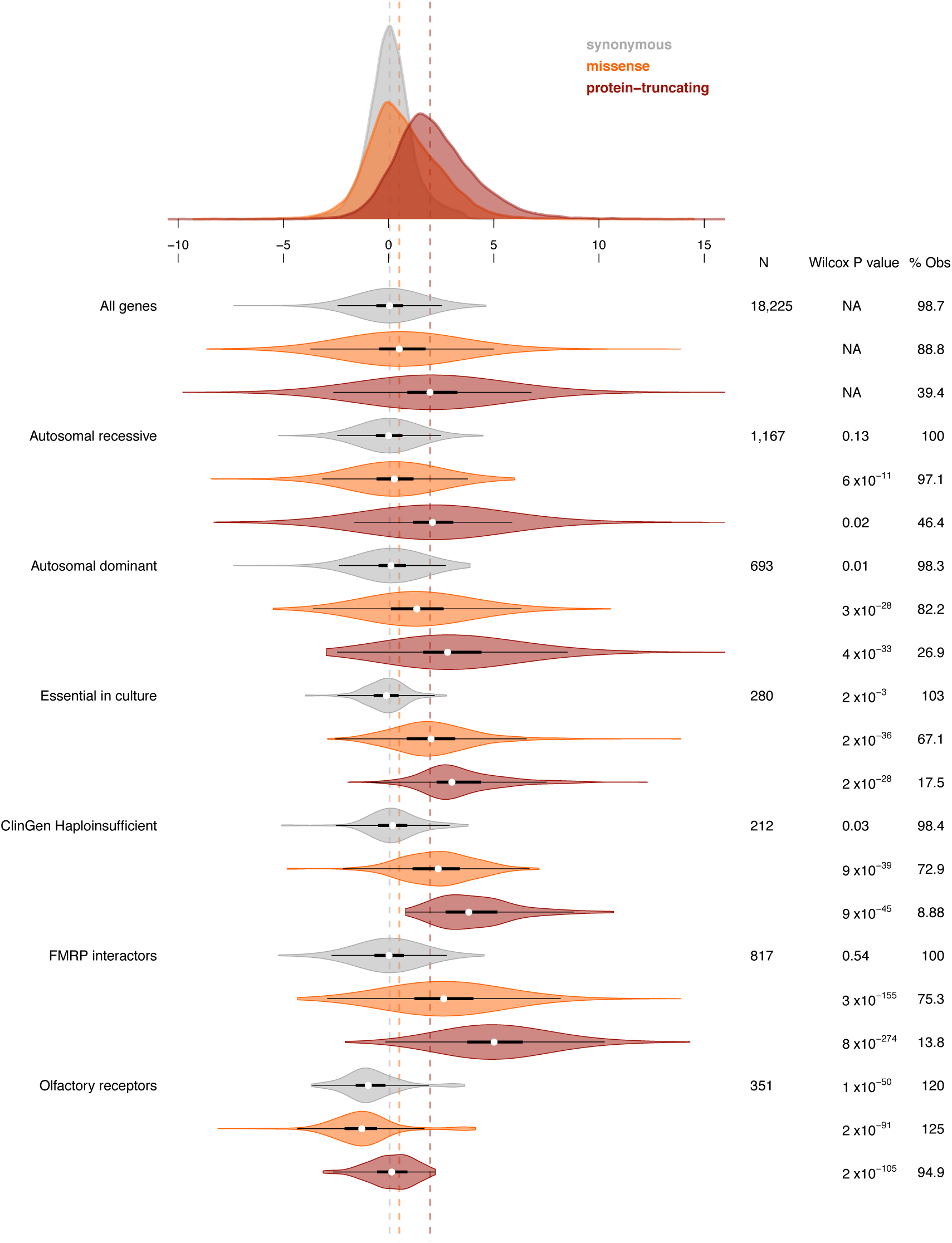
Distribution of synonymous, missense, and protein-truncating Z scores for gene sets. The number of genes in the set, the Wilcoxon p-value for the difference from the full distribution, and the percentage of expected variation observed are reported on the right. Thick black bars indicate the first to third quartiles, with the white circle marking the median.

**Extended Data Figure 7.**
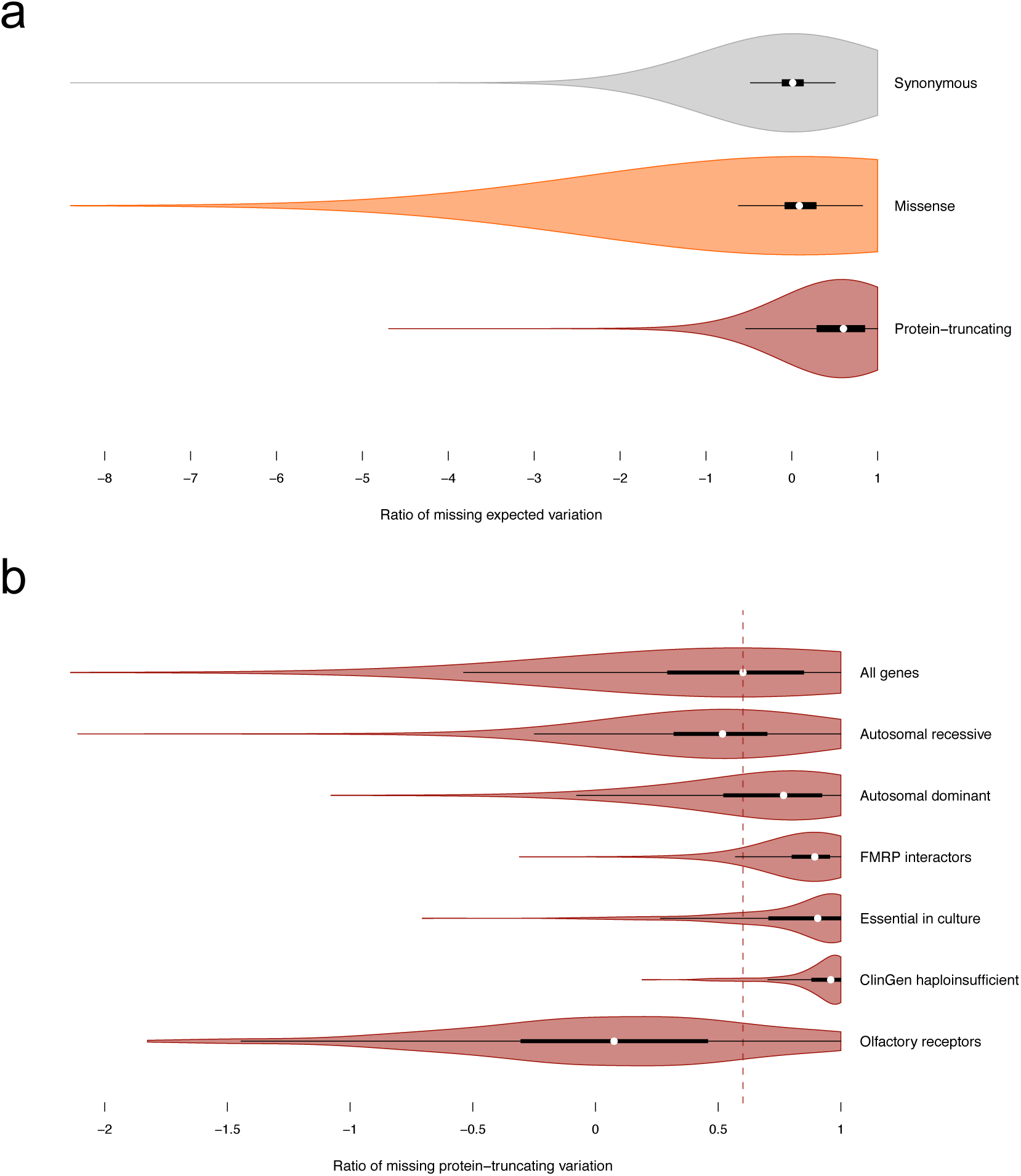
Ratio of missing synonymous, missense and protein truncating variation. a) The distribution of the ratio of missing expected variation for synonymous, missense, and protein-truncating as well as for gene sets of interest. Note that 1 means there were no variants observed and negative values indicate more variation observed than expected. b) The median ratio of missing protein-truncating variation for all transcripts is indicated by the dashed maroon line. For a, the x-axis has been trimmed at −8 (out of −18) to highlight the patterns of the data. Similarly, x-axis in the bottom panel has been trimmed at −2 (out of −5) to highlight the patterns of the data. Thick black bars indicate the first to third quartiles, with the white circle marking the median.

**Extended Data Figure 8.**
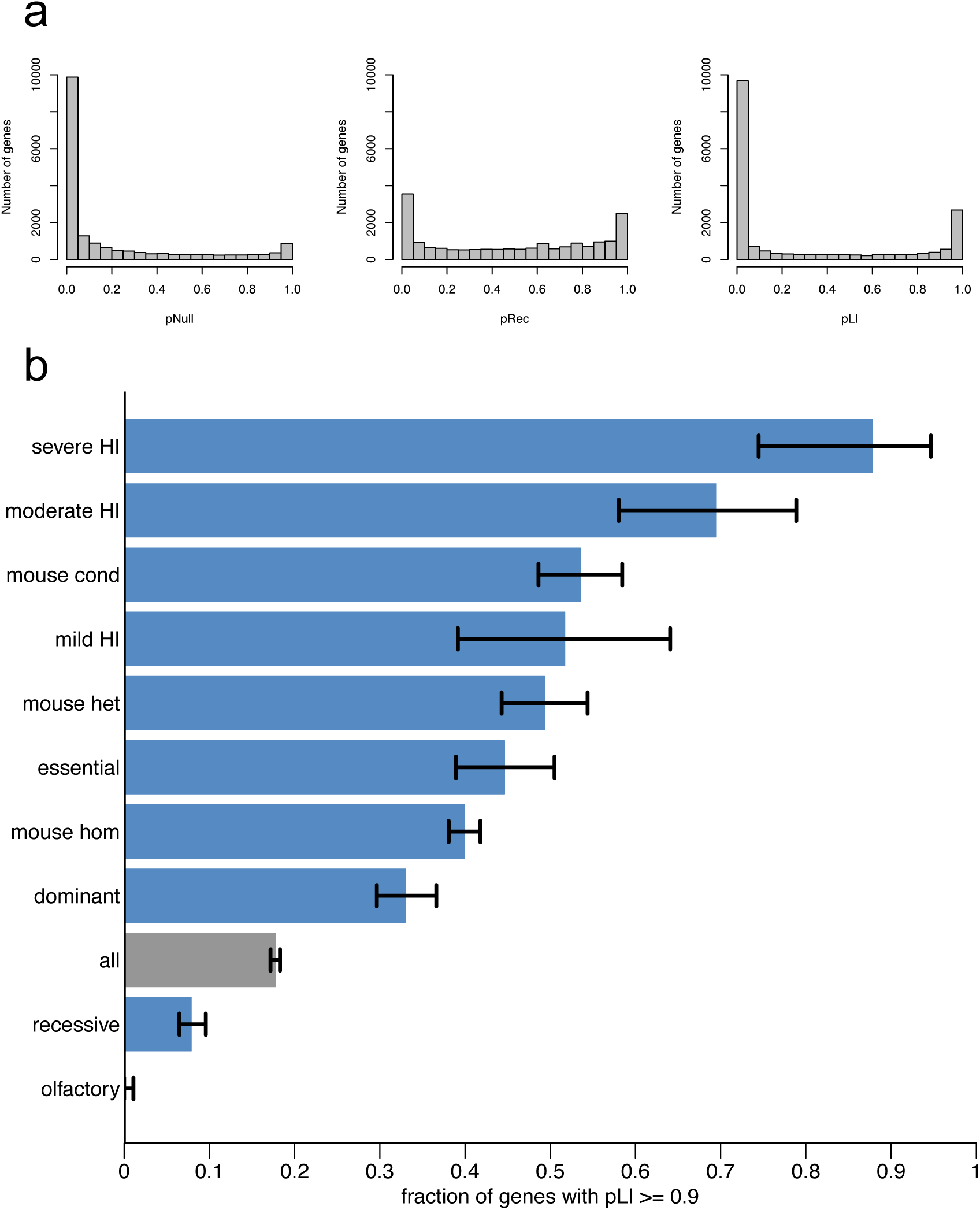
The distribution of pNull, pRec, and pLI across all transcripts and the fraction of genes in a gene set with pLI ≥ 0.9. a) The distributions of pNull, pRec, and pLI for all canonical transcripts. The distribution is roughly bimodal for each. pLI close to one indicates extreme intolerance to loss-of-function variation; we therefore take pLI ≥ 0.9 as the cut-off for extreme loss-of-function intolerance and depict, in b, the fraction of genes from gene sets of interest that have pLI ≥ 0.9. The black error bars indicate a 95% confidence interval. olfactory = olfactory receptor genes (n=371); recessive = recessive disease genes from Blekhman and Berg (n=1,183); all (n=18,225); dominant = dominant disease genes from Blekhman and Berg (n=709); mouse hom = genes that are lethal in mice when both copies are knocked out (n=2,760); essential = genes that are essential in cell culture as curated by Hart et al 2014 (n=285); mouse het = genes that are lethal in mice when one copy is knocked out (n=387); mild HI = haploinsufficient genes that cause a mild disease (n=59); mouse cond = genes that are lethal in mice when conditionally knocked out in adult mice (n=402); moderate HI = haploinsufficient genes that cause moderately severe disease (n=77); severe HI = haploinsufficient genes that cause severe disease (n=44). Please refer to Supplementary Table for more details on gene lists.

**Extended Data Figure 9.**
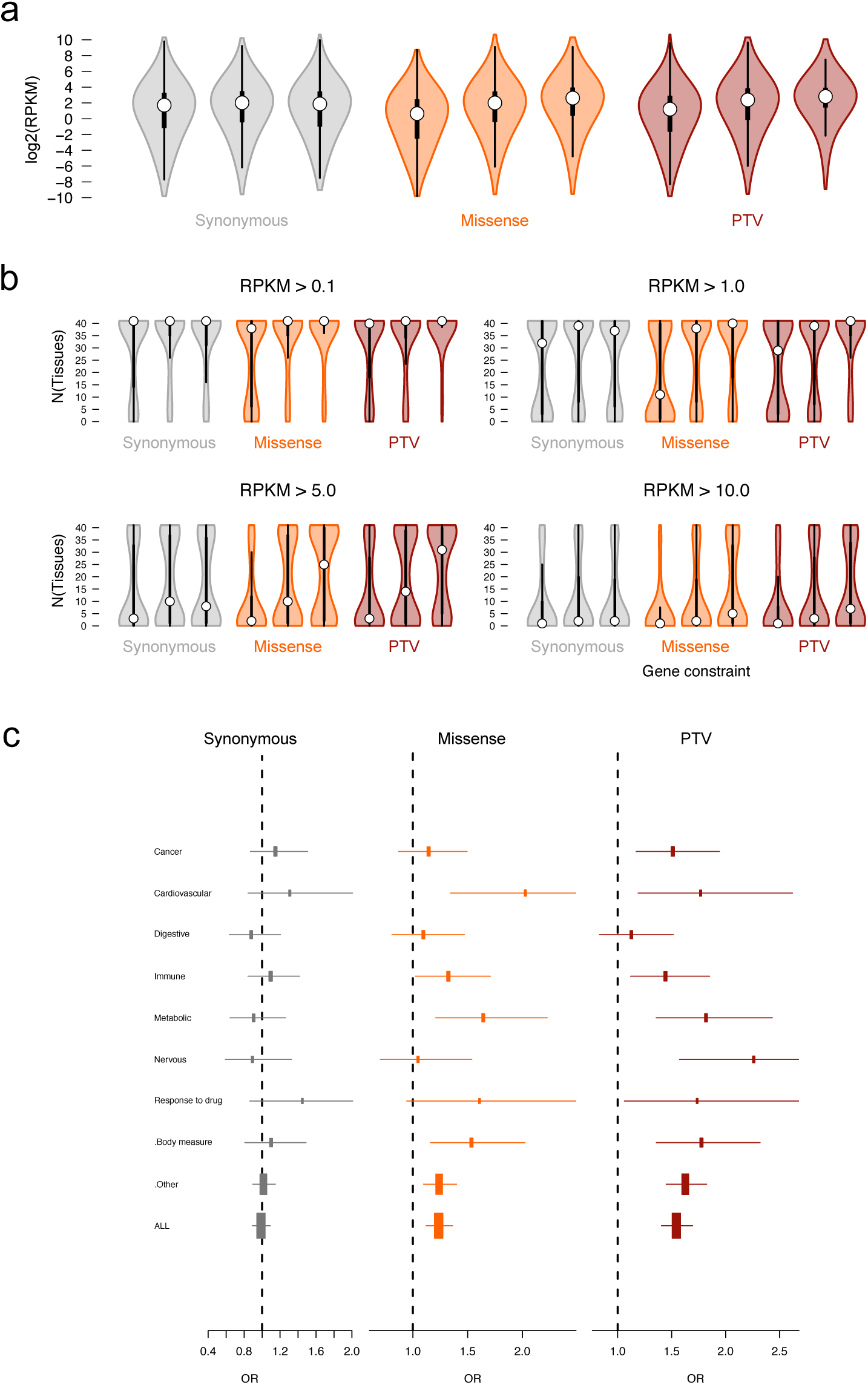
Application of pLI on RNA-Seq data and GWAS hits. a) The relationship between constraint and median gene expression across all tissues. b) The relationship between constraint and tissue expression at different RPKM cutoffs. Thick black bars indicate the first to third quartiles, with the white circle marking the median. c) The odds ratio of being a GWAS hit for each Experimental Factor Ontology trait for the most constrained genes vs the middle bin. The error bars indicate a 95% confidence interval.

**Extended Data Figure 10.**
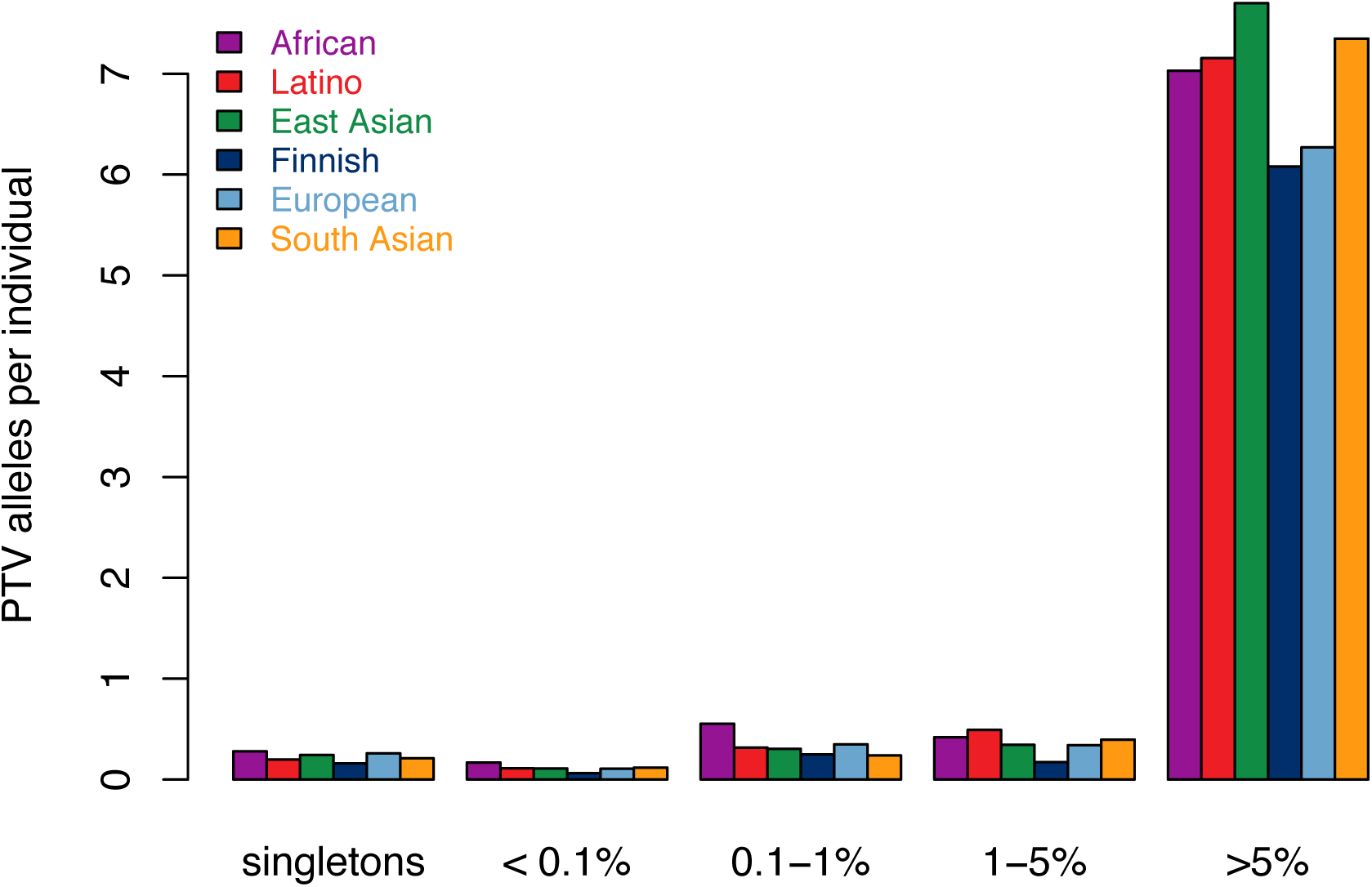
Number of protein-truncating variants in constrained genes per individual by allele frequency bin. Equivalent to Figure 5b limited to constrained (pLI ≥ 0.9) genes.

